# Gaze-centered gating and reactivation of value encoding in orbitofrontal cortex

**DOI:** 10.1101/2023.04.20.537677

**Authors:** Demetrio Ferro, Tyler Cash-Padgett, Maya Zhe Wang, Benjamin Hayden, Rubén Moreno-Bote

**Affiliations:** Center for Brain and Cognition, Universitat Pompeu Fabra, 08002, Barcelona, ES; Department of Information and Communication Technologies, Universitat Pompeu Fabra, 08002, Barcelona, ES; Department of Neuroscience, Center for Magnetic Resonance Research, University of Minnesota, Minneapolis, MN55455, USA

## Abstract

During economic choice, we often consider options in alternation, until we commit to one. Nonetheless, neuroeconomics typically ignores the dynamic aspects of deliberation. We trained macaques to perform a value-based decision-making task where two risky offers were presented in sequence at different locations of the visual field, each followed by a delay epoch where offers were invisible. Subjects looked at the offers in sequence, as expected. Surprisingly, during the delay epochs, we found that subjects still tend to look at empty locations where the visual offers were formerly presented; and, moreover, longer fixation to given empty location increases the probability of choosing the associated offer, even after controlling for the offer values. We show that activity in orbitofrontal cortex (OFC) reflects the value of the gazed offer, but also the value of the offer associated to the gazed spatial location, even if it is not the most recently viewed. This reactivation reflects a reevaluation process, as fluctuations in neural spiking during offer stimuli presentation and delays correlate with upcoming choice. Our results suggest that look-at-nothing gazing triggers the reactivation of a previously seen offer for further reevaluation, revealing novel aspects of deliberation.

## Introduction

Introspection and a myriad of studies imply that eye fixations play important roles in decision making. For instance, when confronted with multiple visual options, we tend to look at them in sequence until we commit to a choice (Horstmann et al., 2009; Orquin & Mueller Loose, 2013; Reutskaja et al., 2011; Russo & Rosen, 1975). In addition, we tend to look longer and most imminently to the offers that we finally choose (Chandon et al., 2009; Krajbich et al., 2010; Russo & Rosen, 1975). This result raises the possibility that there is a causal interplay between eye movements and choice deliberation (Shimojo et al., 2003). At a first glance, the sequential nature of fixations can be thought of as an inevitable consequence of the design fo the visual system, as visual objects need to fall inside the fovea to be accurately processed (Kandel et al., 2000). However, evaluation of options seems to follow a sequential structure also in non-visually guided, abstract decisions (Simon, 1997). Therefore, a sequential evaluation of options may be the default computation in economic choices (Hayden & Moreno-Bote, 2018). This view is supported by the computational benefits of focused evaluation of fewer options as compared to an overall screening of all available options (Mastrogiuseppe & Moreno-Bote, 2022; Meyer et al., 1997; Moreno-Bote et al., 2020) as well as by anatomical and physiological considerations (Rushworth et al., 2011; Yoo & Hayden, 2018).

Neurons in several brain areas, most notably the orbitofrontal cortex (OFC) and ventromedial prefrontal cortex (vmPFC), show modulations in firing rate activity that correlate with the economic value of visual offers (Murray et al., 2007; Padoa-Schioppa, 2007, 2011; Padoa-Schioppa & Assad, 2006; Padoa-Schioppa & Conen, 2017; Rangel et al., 2008; Rich & Wallis, 2016; Rushworth et al., 2011; Schuck et al., 2016; Strait et al., 2014; Wallis, 2007; Wilson et al., 2014; Yoo & Hayden, 2020). One current view is that partially segregated neural populations are selective for different offers in the world, and each independently compute its value as they are compared (Ballesta & Padoa-Schioppa, 2019; Shi et al., 2022). A similar view holds in perceptual decision making, where it has been shown that several competing choices can be prepared in parallel by partially segregated neural populations in premotor and prefrontal areas (Churchland et al., 2008; Cisek & Kalaska, 2010). However, some brain areas represent only the options that are attended to (Butler et al., 2021; Hayden & Moreno-Bote, 2018; Xie et al., 2018), and some work shows that OFC neurons represented offer values in alternation, not in parallel (Rich & Wallis, 2016). Therefore, it is still unclear how many alternative offers can be simultaneously encoded, to what extent offer values are simultaneously evaluated, and how this depends on gaze and attention patterns. Answering these questions is important to understand the canonical computations underlying economic choice.

To tackle these questions, one would need a task where encoding of visual offers (e.g., recognition) and their evaluation are not conflated, which automatically excludes tasks where offers are permanently visible. This is because otherwise it is not possible to determine whether sequential processing of offers is the result of the need of moving the eyes or directing attention in sequence to the visual stimuli. An effective decoupling of encoding and evaluation can be achieved by interleaving the offer presentations with empty screen delay epochs, which is the approach that we adopt here. Further, to test whether evaluation is sequential by default, we let subjects to move their eyes at will, and expect tracking the internal deliberation process by using the look-at-nothing effect (Altmann, 2004; Johansson & Johansson, 2014; Theeuwes et al., 2011), a visuo-motor tendency to gaze to locations currently empty where relevant stimuli were formerly presented. Looking at empty locations is thought to aid memory recollection of the formerly presented visual stimuli (Fourtassi et al., 2014; Johansson & Johansson, 2014) –like looking at the center of the yard to remember the maple tree that used to be there. We hypothesized that it may be associated with memory for choice options and moreover, that it may be associated with active and ongoing evaluation processes that help determine which option is chosen.

We used a two-alternative value-based decision-making task where sequentially presented visual offers left an empty screen after a short presentation. We tracked the free view gaze of two macaques both during offers presentation (look-at-something) and later delay times when screen is empty (look-at-nothing). As expected, we found that animals directed their gaze to the offers, with longer fixations to one offer predicting a higher probability of choosing said offer. The same pattern is observed when the offers become invisible during the delay epoch: the more the empty side is looked at, the higher the probability that the formerly presented offer in that location is subsequently chosen. We confirmed that this effect is independent of the value of the offers by factoring them out in the analysis. In simultaneous recordings of OFC neurons, we confirm the well-known finding that neural activity represents the value of the offer gazed by the subjects. And, more interestingly, looking at the empty location where a previous offer was presented elicits a neural reactivation of its value, regardless of whether another offer was seen and encoded in the interim. This representation reflects an active reevaluation process, as fluctuations of neural activity during look-at-nothing, while controlling for offer values, correlate with choice. We find that the encoding of value during stimulus presentation and reactivation occurs in overlapping cell populations. Our results show an unexpected connection between gazing to empty spaces, reactivation, and reevaluation, which reveals a sequential default mode underlying economic choice.

## Results

### Performance and look-at-nothing gazing in a gambling task

We combine the analysis of choices, oculomotor data and spiking neural recordings of two macaque monkeys performing a visually cued, two-alternative reward gambling task (Fig. 1A; Supp. Tables ST1-ST2 for exact number of trials) (Maisson et al., 2021; Strait et al., 2014). The task consists of the sequential presentation of two vertical bar stimuli (*offer 1* / *offer 2* for 400 ms) at opposite sides of a visual screen display. Each stimulus presentation is followed by an empty screen delay time (*delay 1* / *delay 2* for 600 ms). Subjects are left free to move their gaze at their will during both offer and delay epochs. At the end of *delay 2*, subjects must re-acquire fixation (*re-fixate*) for at least 100 ms at the center of the screen. Upon fixation reacquisition, the two offers appear side by side on their old locations (*choice-go* cue), which instructs the animals to choose one by performing a saccade to the target offer. The reward magnitude 𝑚 of an offer is indicated by the color of the bottom part of its associated rectangular visual stimulus (gray: small; blue: medium; green: large; see Methods), which include a top red part. The height of the bottom part indicates the (success) probability 𝑝 of obtaining the liquid reward of magnitude 𝑚 (cued by its color) if the offer is chosen. The height of the red part corresponds to the complementary probability 1 − 𝑝 for an unsuccessful outcome. If the gamble is unsuccessful, no reward is given. We define the expected value 𝐸𝑉 = 𝑚𝑝 of an offer as the product of its magnitude 𝑚 and its probability 𝑝, and its risk is defined as the value variance, 𝜎^2^ = 𝑚𝑝(1 − 𝑝). We define subjective value as 𝑆𝑉 = 𝑤_1_𝐸𝑉 + 𝑤_2_𝜎^2^, which we compute for each session via logistic regression of choice (Methods Section 4.4; Supp. Table ST3).

**Figure 1:**
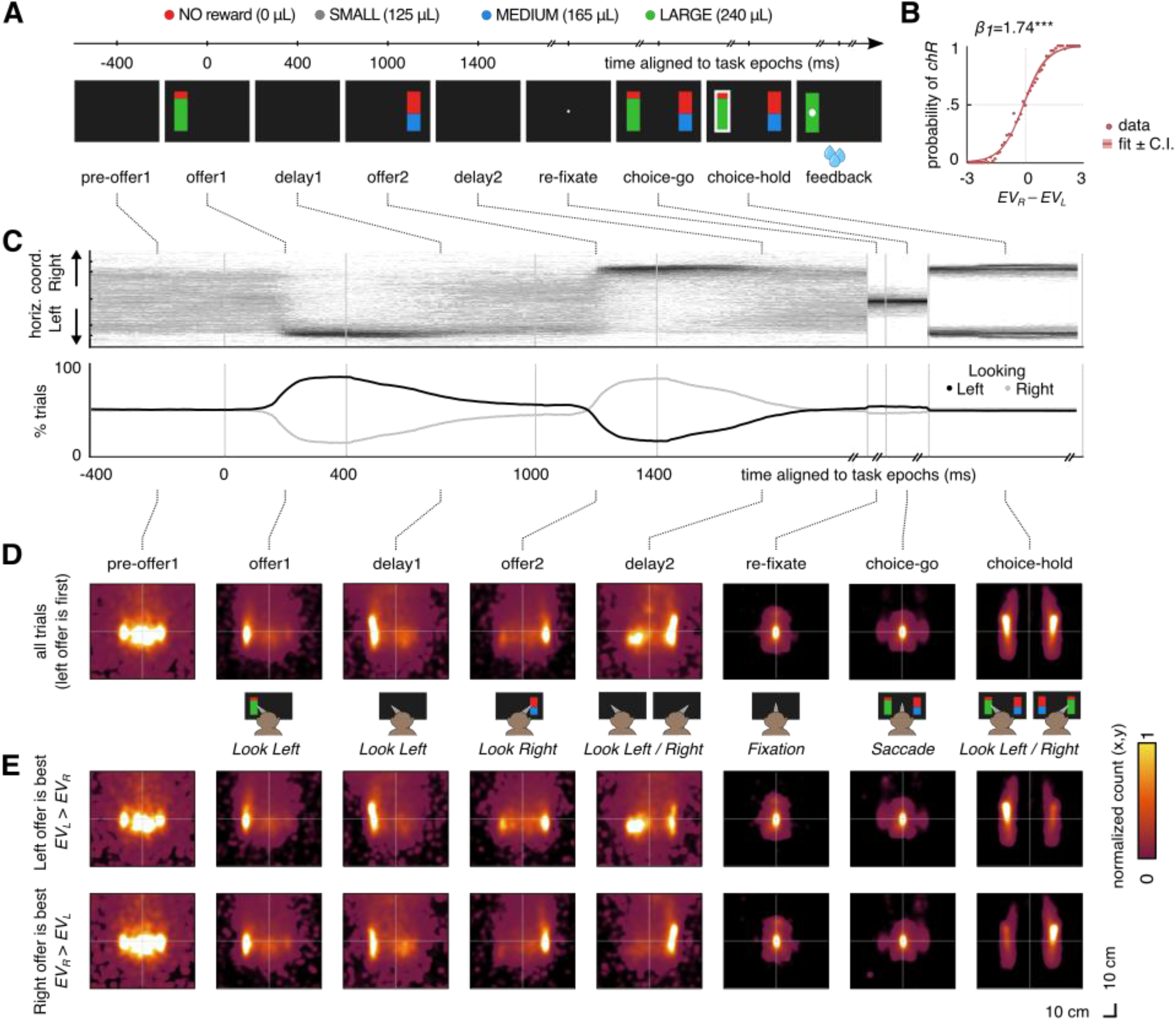
Gambling task and gaze behavior. **A.** Timeline of the two-alternative sequential gambling task for a sample trial. Reward offers are cued by sequential presentation of vertical bar stimuli at opposite screen sides (*offer 1 / offer 2*) for 400ms, interleaved with blank screen delay times (*delay 1 / delay 2*) for 600ms. Stimuli colors either cue a safe (always achieved, reward probability=1), but small fluid reward (gray), or to probabilistic rewards with size medium (blue) or large (green). Reward magnitudes were pseudo-randomized across trials. Reward probabilities of the medium and large offers were independently drawn from a uniform probability distribution. The green or blue portions of the bar is proportional to the reward probability such that if the whole bar is green or blue the probability of the reward would be one. Following fixation to the center of the screen (*re-fixate*), choice is performed after *choice-go* cue by performing a saccade to the selected side. Choice is reported by holding the gaze on the chosen offer for at least 200ms (*choice-hold*). The trial concludes by providing visual feedback and reward or no reward (*feedback*). Reward offers order (first presented Left or Right) were pseudo-randomized to have equal probability across trials. For analysis we mirrored choices and eye data respect to the midline in trials that started with the right offer, so that all trials ‘started’ with the offer located on the ‘left’ screen side. **B.** The probability of choosing the right offer (*chR*) increases with the difference between right and left expected values. Solid line shows logistic fit (logit(*chR*) = 𝛽_0_ + 𝛽_1_(𝐸𝑉_𝑅_ − 𝐸𝑉_𝐿_)); shaded areas show ±95% Confidence Interval (C.I.), *** p<0.001. **C.** Top: distribution of horizontal eye position throughout the trial (spatial bin size is 1 mm, time bin size 1 ms); bottom: fraction (%) of trials separated by looking side (left: average horizontal eye position <0; right: average >0) in 10ms bins during task execution. **D.** Gaze distribution over the visual display throughout the trial. First, subjects preferentially look at the left side, where the offer is first presented (data in trials where the first offer is presented on the right side are mirrored right to left), then they mostly look at the right side when the second offer is presented. Importantly, during the second delay epoch, subjects split gazing to both sizes of the screen, even when the offers are invisible. **E.** Same as D but separated for trials where left offer has the highest EV (top), and for trials where right offer has the highest EV (bottom). Left / Right offer is the best in 48.67% / 49.28% (Supp. Tables ST1, ST2 report the exact number of trials). **A-E.** Data includes 5971 trials with valid behavioral choice report (2463 for subject 1; 3328 for subject 2, Supp. Tables ST1 and ST2 for exact number of trials for each session).

Both subjects perform the task well, by successfully reporting the choice and by following value-based contingencies (most often choosing the offer with the higher value; subject 1: 72.19%; subject 2: 75.72%). We find a significant relationship between choice and difference in value for the two offers (Fig. 1B, logistic regression, p<0.001; Supp. Fig. S1A, for each subject). As the task consists of the sequential presentation of visual offers on opposite sides of the screen, we expect subjects to perform overt visual search to collect sensory information to estimate the value of the offers. Indeed, the eye position of the two subjects mainly follows the stimuli locations (Fig. 1C, top), when offers are presented. In most trials, subjects first fixate the first offer location (*offer 1*: 78.83% and 81.57% of the trials with first offer on the left and right side, respectively; Supp. Table ST2), then the opposite location during the second offer epoch (*offer 2*: 75.78% and 81.82% when second offer is on the left and right side, respectively; Supp. Table ST2). Analogous results are found by computing the fraction of trials in 10 ms time bins where the animal looks at each side of the screen from the midline (Fig. 1C, bottom). We observe that gazing to the visual offers is most prominent in the last 200 ms of the presentation time, once a saccade towards the target has occurred, and partially persists during the delay epochs (Fig. 1C; saccade delay ≈ 150-200 ms; *offer 1 / offer 2* in Supp. Fig. S1B, S1C). After offer and delay epochs, the animals behave according to task instructions by re-acquiring fixation and performing a saccade to one of the two offers to report the choice (Fig. 1C, *re-fixate* to *choice-hold* epochs).

When analyzing eye data in two dimensions mapped on the screen, we find that gaze is concentrated on the offer locations also along vertical axis, i.e., animals preferentially look at the physical rectangular shapes of the offers (Fig. 1D, e.g., *offer 1* and *2* epochs; Supp Fig. S2A for each subject). Interestingly, we observe that gaze locations during delay times show a look-at-nothing bias: animals look at the location of the formerly presented stimuli (*delay 1* and *2* epochs in Fig. 1D), indicating that even in the absence of stimuli, animals recall the spatial location where the stimulus had been presented with a good level of acuity. This effect is not very surprising during *delay 1*, as gaze most often remains on the same side where the first offers has been presented, but less so during *delay 2*: animals asymmetrically split looking time between the two sides of the screen (see the two bumps in *delay 2*), approximately matching the locations of the two formerly presented offers. Overall, the look-at-nothing gazing to empty regions of the screen approximately matches the location of look-at-something gazing to the visual offer when physically presented. These results are observed regardless of whether the first offer is presented on the left or right (Supp. Fig. S1B, S2B). For this reason, for subsequent analyses, we mirrored data by arbitrarily choosing the first offer to be shown on the left, and the second one on the right, in all results. Besides doubling the data size per condition, folding the visual screen to a single reference (by arbitrary convention, we will use left offer first, right offer second) also simplifies the presentation of the following results.

The patterns of look-at-something (*offer 1* and *offer 2*) and look-at-nothing (*delay 1* and *delay 2*) depend on several factors, including the difference in values of the offers. When splitting trials according to the value of the best offer (Fig. 1E; top: best offer is the first one, on the left by convention; bottom: best offer is the second one, on the right), we observe no substantial difference in gaze position during the *offer 1* and *delay 1* epochs, probably because animals sample the first offer in most trials, while the second offer is still unknown. However, during the *offer 2* and *delay 2* epochs, both look-at-something and look-at-nothing gazing are biased towards the location of the more valuable offer. For instance (Fig. 1E, top panel), when the value of the first offer (located on the left) is larger than that of the second offer (on the right), subjects tend to look more on the left side of the screen during the *delay 2* epoch (see Supp. Fig. S2D for comparison). This important pattern shows that look-at-nothing gaze does not simply reflect a memory of the last seen offer, but rather gaze typically reallocates to the location of the visual field that, even if empty, tends to correspond to the better offer. A similar effect is observed during the earlier *offer 2* epoch, for which we find trial occurrences of the gazing over the left side (see bump on the left during the *offer 2* in Fig.1E, top panel), indicating that in some trials subjects barely look at the right side because the first offer has higher value than the second one. The opposite tendency is observed when the expected value of the second offer is higher: subjects mostly look at the right site during the *delay 2* epoch (*offer 2* in Fig.1E, bottom panel). Similar conclusions can be drawn by considering the chosen offer (Supp. Fig. S2C, S2E).

### Gaze position modulates choice during look-at-something and look-at-nothing

To determine the role of gazing in evaluation and choice, we used several analyses to test whether look-at-something and look-at-nothing gazing are predictive of choices. First, we analyzed the relationship between choice and expected value for three overall gazing conditions and for each task epoch: using all trials, or using only trials where the subject spends a larger fraction of time looking at the right side of the screen in that epoch (𝑓_𝑅_ > 0.5), or using trials where the subject spends a larger fraction of time looking the left side of screen (𝑓_𝑅_ < 0.5) (Methods Section 2.3). As shown above (Fig. 1B), the probability of choosing the right offer increases with the value difference 𝐸𝑉_𝑅_ − 𝐸𝑉_𝐿_ (Fig. 2A), but it is also modulated by which side is mostly looked at (mostly right 𝑓_𝑅_ > 0.5 mostly left 𝑓_𝑅_ < 0.5) in all task epochs. Not surprisingly, during the choice-hold epoch, the effect is reflected by a large horizontal shift of the psychometric curve: if 𝑓_𝑅_ > 0.5, the psychometric curves are largely shifted to the left, indicating that the choice is almost always the right (second) offer. But, surprisingly, a significant shift of the curves is also observed in all other task epochs (significance assessment in Supp. Table ST4), being most prominent during *delay 2*. These results show that, after controlling for expected value differences, looking time during both look-and-something and look-at-nothing is predictive of the choice that animal will make at the end of the trial.

**Figure 2.**
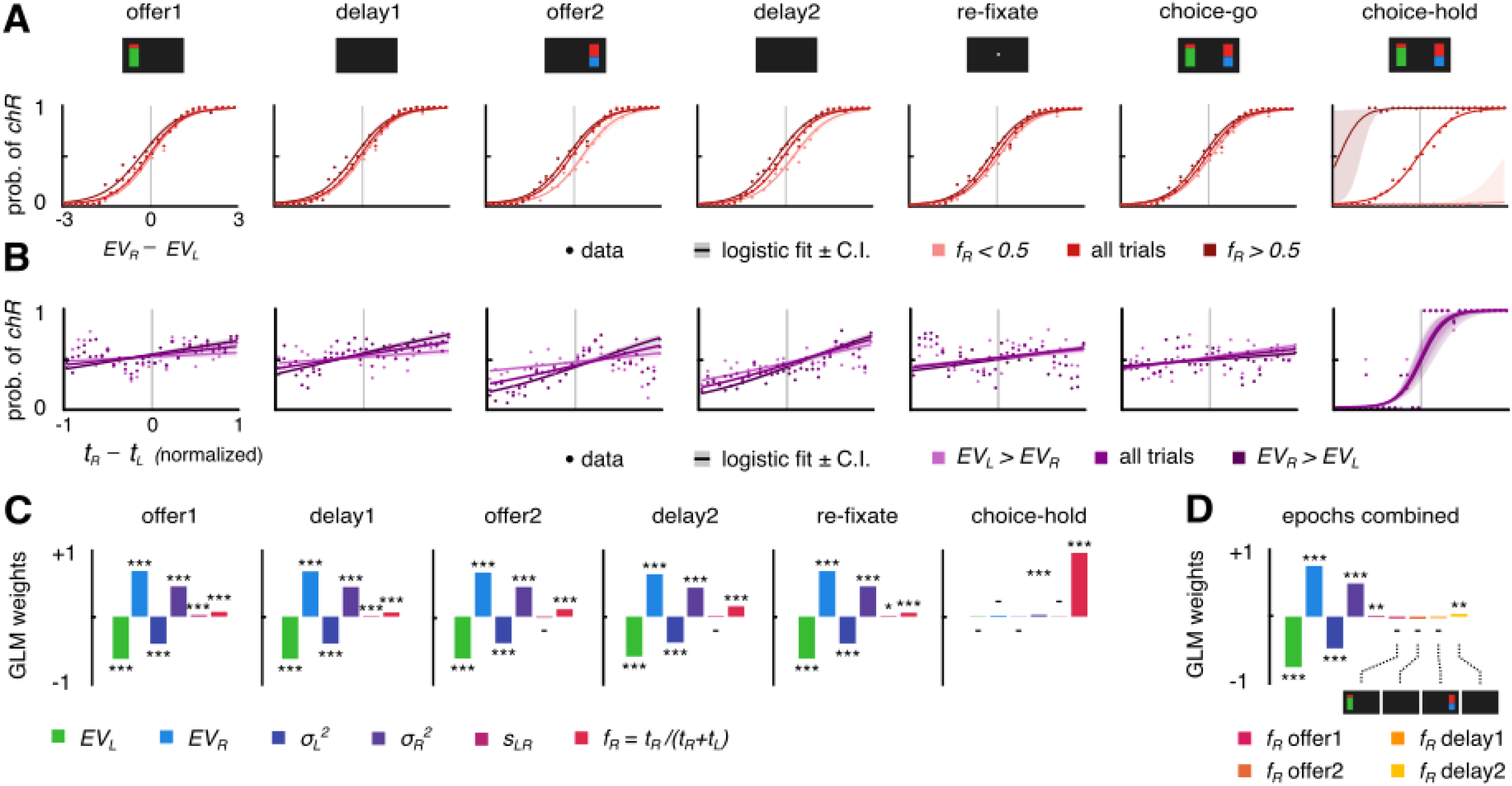
The fraction of time spent looking at either screen side predicts choice, even after controlling for expected value and risks of the offers. **A.** Probability of choosing the right offer as a function of difference in EV of the two offers 𝐸𝑉_𝑅_ − 𝐸𝑉_𝐿_. Solid lines are logistic regression fits logit(𝑐ℎ𝑅) = 𝛽_0_ + 𝛽_1_(𝐸𝑉_𝑅_ − 𝐸𝑉_𝐿_), shaded areas are ±95% C.I. The analysis is applied in each epoch for all trials where subjects mostly look to the left (𝑓_𝑅_ = 𝑡_𝑅_/(𝑡_𝑅_ + 𝑡_𝐿_) < 0.5) or to the right (𝑓_𝑅_ > 0.5) screen side. **B.** Probability of choosing the right offer as a function of the difference in the right- and left-screen looking times 𝑡_𝑅_ − 𝑡_𝐿_. Solid lines are logistic regression fits logit(𝑐ℎ𝑅) = 𝛽_0_ + 𝛽_1_(𝑡_𝑅_ − 𝑡_𝐿_), shaded areas are ±95% C.I. The analysis is applied in each time window for all trials, for trials with the highest 𝐸𝑉 on the left and for trials with the highest 𝐸𝑉 on the right. **C.** Logistic regression of behavioral choice. Regressors include the expected value of left and right offers 𝐸𝑉_𝐿_, 𝐸𝑉_𝑅_, variance (risk) of the two offers 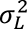, 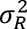, order of presentation (𝑠_𝐿𝑅_ = +1 if first offer is on the left; −1 if it is on the right), and fraction of time looking at right screen side 𝑓_𝑅_. **D.** Logistic regression model of regression of behavioral choice, using all regressors used in C along with the fractions of right-looking time 𝑓_𝑅_ for most relevant task epochs (*offer 1, delay 1, offer 2, delay 2*). **A-D.** Data include 𝑛 = 5971 trials with valid behavioral choice report (𝑛 = 2463 for subject 1; 𝑛 = 3328 for subject 2, Supp. Tables ST1 and ST2 for exact number of trials). Pooling across trials is made with reference to the first offer side: data in trials with first offer on the Right side are horizontally mirrored. Significance levels: *p<0.05, **p<0.01, ***p<0.001.

In a second analysis, we find that the effect of looking time on the probability of choice is graded (Fig. 2B). When plotting the probability of choosing the right offer as a function of the difference of the time looking on the right minus the time looking on the left side (𝑡_𝑅_ − 𝑡_𝐿_), we find that it monotonically increases for all task epochs, regardless of the expected value difference (all trials). The same monotonic increase is observed when dividing trials by the expected offers: higher for left offer (𝐸𝑉_𝐿_ > 𝐸𝑉_𝑅_), and higher for right offer (𝐸𝑉_𝑅_ > 𝐸𝑉_𝐿_). Note that the slope is larger when the right offer value is higher as we analyze the probability of choosing the right offer (Supp. Table ST5).

To predict choice on a trial-by-trial basis, we used a logistic regression model that included as regressors the values of the offers, their risk, the location of the first offer, and the fraction of time that the animal looks at the right side each epoch (Fig. 2C; Supp. Fig. S3A for each animal; Methods Section 2.3). Adding both the expected value of the offer and their risk is important to control for other factors known to determine choices and could affect gaze position. We find, as expected, that in all epochs from *offer 1* to *re-fixate* epochs, the value of the offers are the strongest regressors (Fig. 2C). The risk of each offer (see Methods) has also a positive contribution to the choice probability, in accordance with the well-known risk-seeking behavior of monkeys (Heilbronner & Hayden, 2013). In the *choice-hold* epoch, the strongest regressor is the fraction of looking time, due to the causal role of visual fixation and choice by task construction. Confirming the previous analyses, in earlier epochs (*offer 1* to *re-fixate*), the fraction of looking at the right side has a small but significant effect on the final choice. This effect is not only observed during the offer epochs, but also in the delay epochs, when no stimulus is displayed. Thus, in the absence of stimuli, the animals have an increased probability of choosing the right offer if they look at the right empty side of the screen for a longer duration, and the opposite when looking more to the left empty side, even after controlling for the expected values and risks of each offer. Importantly, the results are reproduced when using for the analysis the fraction of time that animals spend looking over the rectangles that define the location of the offers (Supp. Fig. S3B). We also asked whether the gaze patterns during the blank epochs can account by themselves for some variability in choice even after accounting for gaze during the stimulus epochs. We therefore used a new logistic regression model where the fractions of looking at the right side for all epochs prior to choice are included as regressors (*offer 1 / 2, delay 1 / 2,* Fig. 2D). This regression again shows that the fraction of time looking at the right during the *delay 2* epoch has a significant effect on choice (p<0.01, Fig. 2D).

### Look-at-something and look-at-nothing value encoding

While look-at-nothing gazing is suggestive of an evaluation process that persists after stimulus disappearance from the visual field and contributes to the choice, to further support this hypothesis we need to show that neural activity indeed reflects re-evaluation of the offers. We recorded responses from n=248 neurons (𝑛 = 163 for subject 1, 𝑛 = 85 for subject 2, Supp. Table ST1) in two core reward regions, areas 11 and 13 of orbitofrontal cortex OFC (Fig. 3A). The importance of OFC and reward value estimation is well established (Murray et al., 2007; Padoa-Schioppa, 2011; Padoa-Schioppa & Conen, 2017; Rushworth et al., 2011; Schuck et al., 2016; Wallis, 2007; Wilson et al., 2014). We first studied the neural encoding of expected value of the offers by analyzing the evoked spiking activity at different task epochs. To do so, we compute the average number of spikes in sliding time windows of 200 ms, shifted by 10 ms steps, for each trial (𝑛 = 5791 total trials, 𝑛 = 2643 for Subject 1; 𝑛 = 3328 for Subject 2; Supp. Tables ST1-ST2 for details). To calculate the fraction of neurons that encode the value of each of the two offers, we used linear regression models where the activity of each neuron was exclusively predicted by the 𝐸𝑉 of either offer (first or second, respectively: left 𝐸𝑉_𝐿_, or right 𝐸𝑉_𝑅_) and computed the fraction of neurons with significant regression weights (Methods Section 4.1; significances is assessed both from *F* and permutation tests). To control for spurious fluctuations in the significance of the fraction of significant neurons in short sequences of times, we performed a cluster-based run-length analysis (Butler et al., 2021; Methods Section 4.1).

**Figure 3.**
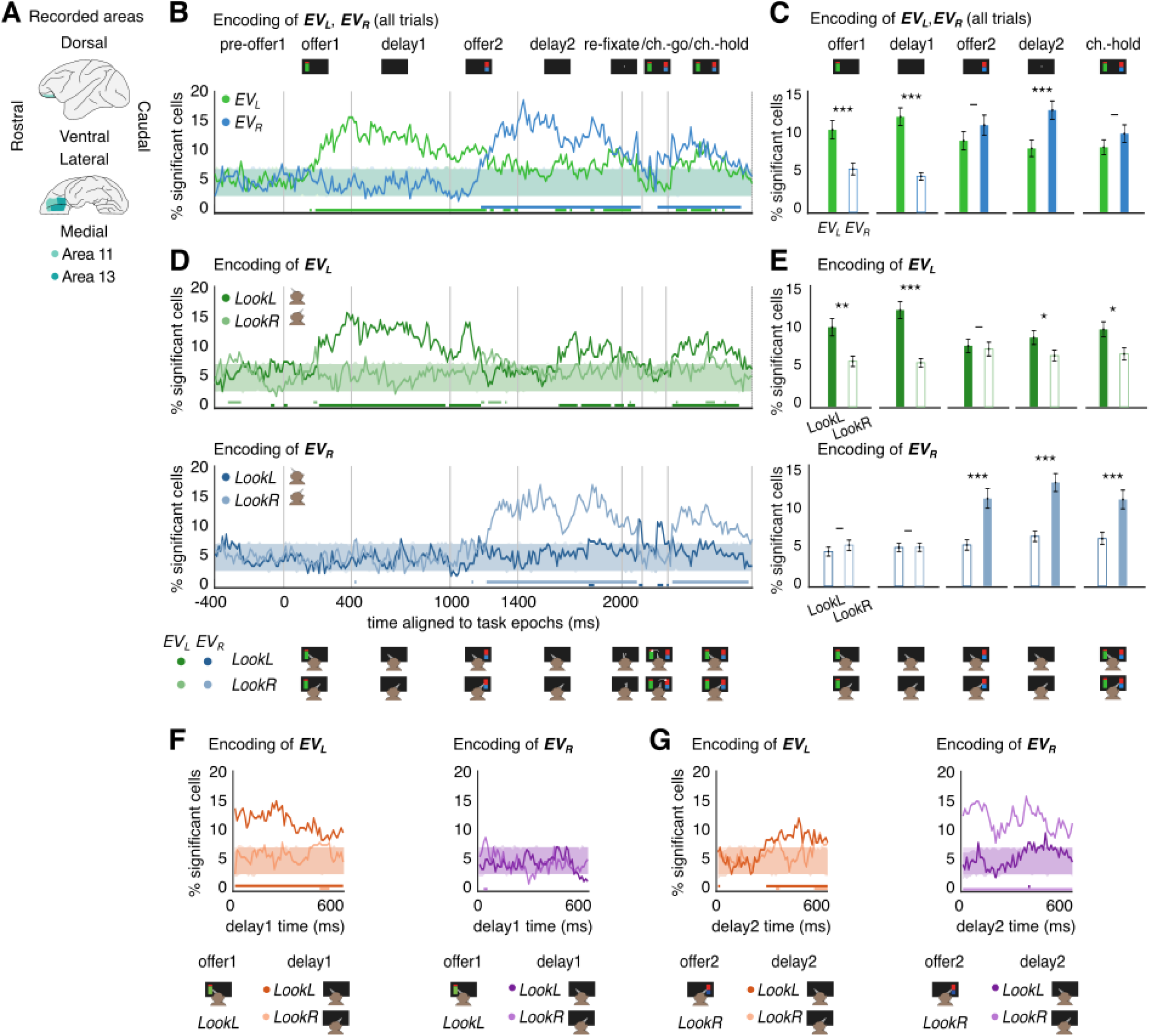
Gaze-dependent encoding of the EV of the offers and reactivation in OFC. **A.** Recorded brain areas, covering macaque brain areas 11 and 13 in OFC. **B.** Fraction of cells showing significant encoding of offer expected value (either 𝐸𝑉_𝐿_ or 𝐸𝑉_𝑅_) for each neuron using the linear models 𝜂 = 𝛽_0_ + 𝛽_1_𝐸𝑉_𝐿_ (green) and 𝜂 = 𝛽_0_ + 𝛽_1_𝐸𝑉_𝑅_ (blue) where 𝜂 is the spike rate of that neuron in each trial. The regression using all trials, regardless of where animals are looking at. Solid lines show the empirical fraction in each time 10ms bin; shaded areas (overlaid, green and blue, in opacity) cover the 5^th^ to 95^th^ percentile of significant fraction of cells encoding the respective 𝐸𝑉 obtained via 𝑛 = 1000 trial order permutations; bottom lines report consecutive runs of time bins with significant encoding of 𝐸𝑉_𝐿_ or 𝐸𝑉_𝑅_ (assessed via run length analysis, detecting significantly long bursts with fraction of encoding cells above the 95^th^ percentile of trial-order shuffled data). **C.** Time-averaged fraction of cells showing significant encoding of offer 𝐸𝑉s. The significance of empirical values is assessed as exceeding of the 95^th^ percentile of the same values run over 𝑛 = 1000 trial-order shuffles of the data (non-significant bars are reported in white, with colored frame). Results for the fraction of cells for 𝐸𝑉_𝐿_ and 𝐸𝑉_𝑅_ are compared via non-parametric tests (two-sided Wilcoxon signed-rank test), significance: - n.s., *p<0.01, **p<0.01, ***p<0.001. **D.** Top: fraction of significant cells encoding 𝐸𝑉_𝐿_ using trials where subjects mostly *LookL* (average eye position for current bin <0, green) with trials where subjects mostly *LookR* (average eye position >0, light green). Bottom: same as before but focusing on 𝐸𝑉_𝑅_ and comparing *LookL* trials (blue) with *LookR* trials (light blue). **E.** Top: same as C but focusing on 𝐸𝑉_𝐿_ and comparing *LookL* with *LookR* trials. Bottom: same as C but focusing on 𝐸𝑉_𝑅_ and comparing *LookL* with *LookR* trials. Comparisons of looking sides are run via non-parametric (one-sided Wilcoxon signed-rank tests), testing that directing gaze to either side yields stronger encoding for ipsilateral 𝐸𝑉. **F.** Encoding of offer 𝐸𝑉s at *delay 1* time for trials when subjects gazed to left side of the screen during the last 200 ms of *offer 1* presentation. Results show the encoding of 𝐸𝑉_𝐿_ (left) and 𝐸𝑉_𝑅_ (right), comparing *LookL* (orange for 𝐸𝑉_𝐿_, purple for 𝐸𝑉_𝑅_) and *LookR* (light orange for 𝐸𝑉_𝐿_, light purple for 𝐸𝑉_𝑅_). Solid lines, shaded areas and bottom lines are computed as in B. **G.** Same as F, but focusing on *delay 2* time, for trials when subjects inspected the right side of the screen during the last 200 ms of *offer 2* presentation. **F-G.** Complementary cases, i.e., comparisons between *LookL* and *LookR* at *delay 1* for *LookR* during the last 200 ms of *offer 1*, and comparisons between *LookL* and *LookR* at *delay 2* for *LookL* during the last 200 ms of *offer 2,* are shown in Supp. Fig. S11. **B-G.** Monkeys and screen display icons are reported to support the visualization by recalling task conditions and respective main gaze patterns during task epochs.

First, using all the trials (without conditioning on gaze), we find that the fraction of neurons encoding either the first or second expected values is not significantly different from chance in the *pre-offer 1* epoch time, as expected, but it is strongly modulated in the following epochs in a sensible way (Fig. 3B). Specifically, the fraction of neurons that encode the value of the first offer (Fig. 3B, 𝐸𝑉_𝐿_, green line) increases significantly above chance (shaded area) in the *offer 1* epoch after a latency of 150-200 ms, and it remains above chance for almost all *delay 1* time. In contrast, the fraction of neurons that encode the value of the second offer (𝐸𝑉_𝑅_, blue line), is not significantly above chance during both *offer 1* and *delay 1* epochs, as it has not been presented yet. During the *offer 2* and *delay 1* epochs the reversed pattern is observed: there is a significant encoding of the value of the second offer (blue line), while the encoding of the first value seems to vanish. The absence of encoding of the expected value of the first offer during the *offer 2* epoch is consistent with the idea that in most trials the subjects look at the location of the second offer, and therefore encoding of the second offer grows in alternation to the encoding of the first one. The difference in encoding at each task epoch of the expected values of the first and second offers is also evident when comparing the fractions of neurons encoding each value averaged across task epochs (Fig. 3C).

We next asked whether gaze direction modulates neural encoding, even in epochs where the screen is empty. For each 10 ms bin in each trial, we separate trials depending on whether animals mainly look at the right or left sides of the screen, based on whether the average eye position in that bin is negative or positive, respectively (Methods Section 4.2). We first corroborate what has been observed during look-at-something gazing: in trials and bins where animals look at the left side of the screen (*LookL*), we find that there is a significant fraction of neurons encoding the first offer during the *offer 1* epoch (Fig. 3D top, *LookL*, dark green line), with magnitude and temporal profile comparable to the gaze-independent encoding of that value (compare with Fig. 3B, green line). Interestingly, in trials where the animal does not look at the first offer side during the *offer 1* epoch (Fig. 3D top, *LookR*, light green line), the encoding of the expected value of the first offer is not present. As expected, the value of the second offer during the *offer 1* epoch is not significant (Fig. 3D bottom) regardless of the average gaze position within the epoch (*LookL*, dark blue line; *LookR*, light blue), as the second offer has not been presented yet.

The same pattern of results described so far is reproduced during the *delay 1* epoch (Fig. 3D, *delay 1*), with the relevant novelty that during this epoch the first offer is no longer visible. Thus, for look-at-nothing gazing we find that *delay 1* activity is significantly modulated by offer value (Fig. 3D top, dark green line). We argue that this encoding is not simply a passive, gaze-independent memory of the formerly presented first offer: we find that by using only the trials where subjects look at the first offer during *offer 1* (*LookL* during the last 200 ms of *offer 1*), but then look at the opposite side during *delay 1* (*LookR* during the last 200 ms of *delay 1*), there is lack of encoding of the expected value of the first offer (Fig. 3F, left panel, light orange; 32.26% of trials, Supp. Table ST2). Thus, looking at the empty side where no previous offer has been presented so far washes out any significant trace of the value of the first seen offer, suggesting that its memory persistence or lack of thereof depends on gaze.

During the *offer 2* epoch, we observe the reversed pattern compared to the *offer 1*: there is significant encoding of the expected value of the second offer when subjects look to the right, where second offer is located, (Fig. 3D bottom, *offer 2,* light blue line) and absence of encoding when the left side is gazed (Fig. 3D bottom, dark blue). Again, during the *delay 2* epoch, where the screen is empty, we find that there is encoding of the second offer only when the animal looks at the right (Fig. 3D bottom, light blue line). Crucially, if the animal looks at the left side during the *delay 2* epoch, there is a significant re-activation of the encoding of the first offer (Fig. 3D top, *delay 2*, dark green line). Therefore, look-at-nothing gazing during the *delay 2* epoch leads to the maintenance of neural representation of the most recently presented (second) offer value when looking at its former location (*LookR*), and to the reactivation of the first offer value if looking at the first offer presentation side (*LookL*). This observation cannot be accounted for by a pure memory trace of the last seen offer coming from trials where the animals simply have not looked at the second offer (which occurs in 21.2% of the trials, Supp. Table ST2): when the second offer is gazed during the *offer 2* epoch, but then gaze is diverted to the opposite side during the *delay 2* epoch (36.48% of the trials, Supp. Table ST2), we find a significant reactivation of the expected value of the first offer (Fig. 3G, left panel, dark orange line). As controls, we observe the maintenance of the encoding of the value of the second offer when the animals keep looking at the right side of the empty screen in the *delay 2* epoch after looking at the second offer in the *offer 2* epoch (Fig. 3G, right; 43.32% of the trials, Supp. Table ST2).

All above results were confirmed comparing the fraction of neurons significantly encoding either the left or right expected values when conditioning on trials where the animals looked on the right or on the left of the screen by considering average fractions of significant cells in each epoch time (Fig. 3E; Methods Section 4.2). We also confirmed that results in Fig. 3 hold for each individual subject (Supp. Fig. S5), using window sizes of 150 ms and 250 ms instead of 200 ms (Supp. Fig. S6), normalizing activity by subtraction of the *pre-offer 1* activity (Supp. Fig. S7, Methods Section 4.1), taking the coefficient of determination 𝑅^2^ of the linear fits as statistics instead of the fraction of cells (Supp. Fig. S8A,C; Methods Section 4.1), and using model-free permutation tests instead of F-test statistics to test significance (Supp. Fig. S8B,D; Methods Section 4.1). Further, since the trial pools for *LookL* and *LookR* have in general a different size, we repeated our analysis, and reproduced the results, by subsampling to pools of even size (Supp. Fig. S9; Methods Section 4.1).

### Reactivation of value encoding during look-at-nothing correlates with choice and is mediated by neural populations overlapping the ones encoding value during look-at-something

A final crucial question is whether activity in OFC during look-at-nothing simply reflects a reactivation of a spatial association of the formerly presented offer, or whether there is further re-evaluation during this reactivation. To answer this question, we performed two analyses. First, we used a linear model to predict the activity of OFC neurons using as regressors the subjective value (𝑆𝑉) of the offers and the choice of the animal at the end of the trial. We argue that if look-at-nothing gazing reflects reevaluation of the offers, we should see in the delay epochs a significant effect of choice that cannot be accounted for by the subjective values of the offers alone. Indeed, in a gaze-independent analysis, a significant fraction of neurons carries choice signals through the last task epochs and became prominent during the choice epoch (Fig. 4A top, black line). Therefore, fluctuations of activity of the neurons are accounted for by for choices beyond what would be predicted by the subjective values of the offers alone, suggesting a role of the OFC activity in the choice, in accordance with previous literature (Ballesta et al., 2020, 2022; Strait et al., 2014). And importantly, similar patterns are revealed when conditioning on gazed location to an offer or its empty location (Fig. 4 middle and bottom), suggesting that evaluation and reevaluation of the offers continues through late task epochs (see Supp. Fig. S11 for a replication of the results using 𝑆𝑉s instead of 𝐸𝑉s).

**Figure 4.**
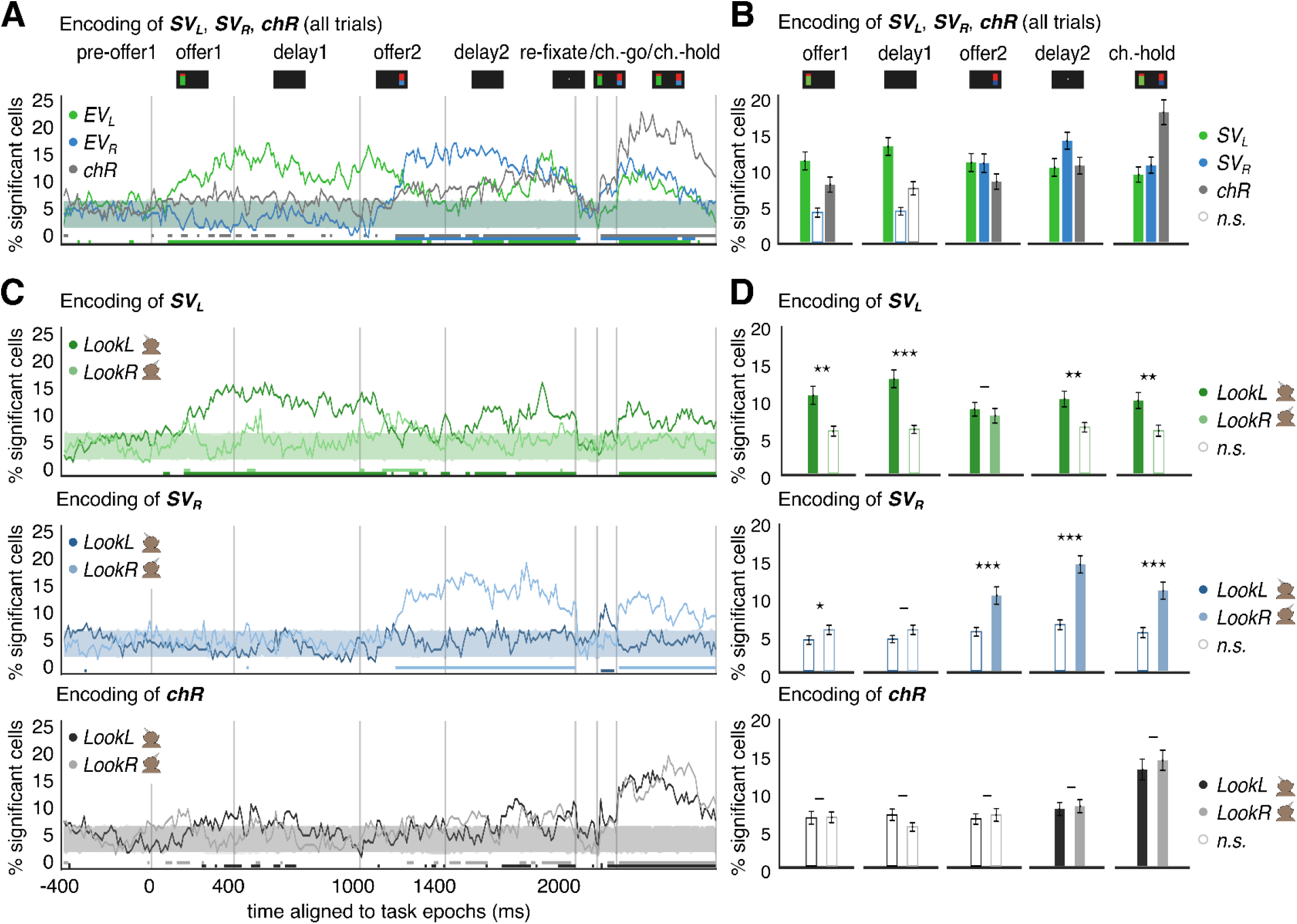
Choice-related activity supports the re-activation of offer value encoding. **A.** Fraction of cells encoding for offer subjective value (𝑆𝑉_𝐿_, 𝑆𝑉_𝑅_) and choice (𝑐ℎ𝑅 = 1 right offer is chosen, 0 otherwise), fit to a linear model of the spike rate 𝜂, as 𝜂 = 𝛽_0_ + 𝛽_1_𝑆𝑉_𝑅_ + 𝛽_2_𝑆𝑉_𝐿_ + 𝛽_3_𝑐ℎ𝑅 using all trials (thus ignoring gaze position). Subjective Values (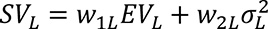 and 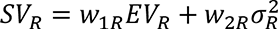; 𝑤_1𝐿_, 𝑤_2𝐿_, 𝑤_1𝑅_, 𝑤_2𝑅_) are computed in each session via regression of the choice: 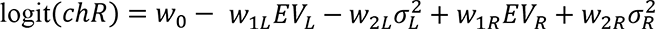 (Supp. Table ST3). Solid lines, shaded areas and bottom lines are computed as in Fig. 3B. **B.** Time-averaged fraction of cells showing significant encoding of offer 𝑆𝑉s and 𝑐ℎ𝑅, computed as in Fig. 3D. **C.** Top: same as A, focusing on the encoding of 𝑆𝑉_𝐿_, comparing trials where subjects mostly *LookL* vs *LookR* in each time bin; middle: same as A, focusing on the encoding of 𝑆𝑉_𝑅_, comparing *LookL* vs *LookR*; bottom: same as A, focusing on the encoding of 𝑐ℎ𝑅, comparing *LookL* vs *LookR*. **D.** Top: same as B but focusing on 𝑆𝑉_𝐿_ and comparing *LookL* vs *LookR;* Middle: same as top, but for 𝑆𝑉_𝑅_; Bottom: same as top, but for 𝑐ℎ𝑅. The time-averaged fraction of time bins with significant encoding for *LookL* and *LookR* are compared via non-parametric tests (Wilcoxon signed rank, one-tailed for 𝑆𝑉s, testing that ipsilateral look yields higher fraction of encoding cells; two-sided for 𝑐ℎ𝑅, testing for non-zero mean in their difference) significance levels: - n.s., *p<0.05, **p<0.01, ***p<0.001.

Second, if reactivation of the formerly presented offer reflects reevaluation, we expect to see that the same neurons participate in both the encoding of the value and in its reactivation. Thus, we considered the encoding of either offer in combination with look-at-something and look-at-nothing. We observe that the encoding weights for the value of an offer when is looked at correlate across neurons with the encoding weights for the value of the same offer when looking at its empty location during *delay 2* (*offer 1*: 𝜌 = 0.49, *offer 2*: 𝜌 = 0.23; *p* < 0.001 in both cases; Supp. Fig. S11A; weights are computed using the last 200 ms of each relevant epoch). This confirms that overlapping neural populations are responsible for both value encoding and value reactivation. However, the same weights are negatively correlated when subjects look to the opposite location during *delay 2* (*offer 1*: 𝜌 = −0.34, *offer 2*: 𝜌 = −0.26; *p* < 0.001 in both cases; Supp. Fig. S11B), suggesting that activity decays during the reactivation of the value of the alternative offer.

## Discussion

Using an economic choice task where offers are sequentially presented and free fixation, we show that gaze can reactivate neural responses that reflect the values of previously shown offers, presumably to facilitate their re-evaluation for the formation of a choice. After accounting for value, residual variability in choice is explained by two additional factors: how much time the subjects were looking at the offers when presented (a look-at-something bias), and, in addition, how much time they look at the locations where they were formerly presented in empty screens (a look-at-nothing bias). Therefore, gaze biases choice regardless of the value of the visual stimulus. Based on single-neuron activity in OFC, we find that (1) gazing to an offer triggers a neural encoding of its value, (2) gazing to an empty region where no stimulus has been presented washes out that value encoding, and (3) gazing to an empty region where a stimulus was formerly presented re-activates its value encoding. Furthermore, variability in the size of this reactivation predicts choice, and overlapping neural population are involved in both sensory encoding of value and its reactivation during look-at-nothing, suggesting a role of re-activation in the formation of the choice. Altogether, our data and results eloquently speak in favor of an active role of gaze in decision making by facilitating memory re-activation and offers reevaluation.

Our study is based on a large body of research showing that gaze is important for decision making (Horstmann et al., 2009; Krajbich et al., 2010; Orquin & Mueller Loose, 2013; Reutskaja et al., 2011; Russo & Rosen, 1975; Shimojo et al., 2003), but to our knowledge ours is the first work that studies look-at-nothing gazing in neural data. Previous works that left stimuli over the screen (Butler et al., 2021; McGinty et al., 2016) also observe a reduction in value encoding as soon as gaze moved away from the physical stimulus, consistent with our results. Our work shows the more intriguing result that looking at regions of space where there was a stimulus triggers the re-activation of its associated memory. This reactivation is not a simple memory trace of the most recently gazed stimulus, as reactivation of the first offer occurs even when fixation is broken by looking to a second offer in the meantime, eventually playing the role of a distractor. Our results are in line with previous theorization of looking-at-nothing gazing, that has proposed that gaze is used as memory spatial indexing mechanism (Platzer et al., 2014; Renkewitz & Jahn, 2012), thus providing neural support for the of gaze-triggered memory recollection hypothesis. Morevoer, these results suggest the reactivation process is itself associated with the evaluation process and is not simply a passive recollection.

Gaze in other conditions has also been shown to re-activate sensory memories. For instance, work in rodents has shown that during rapid eye movement (REM) sleep, eye movements trigger neural activity in the anterodorsal nucleus of the thalamus that reflects internal heading in the virtual world of the sleeping animal, showing a tight coupling between eye and internal variables (Senzai & Scanziani, 2022). Our work in addition shows that the coupling between eye position and internal variables extends to awake primates and might subserve an important mechanism for memory re-activation during decision making. Further, our results extend those of re-activation in the hippocampus for memory reconsolidation (Grella et al., 2022) and planning (Foster & Wilson, 2006), but here we show that it can also be a mechanism for re-evaluation of information in cortex to aid visually guided economic decision making.

We have shown the moving the eyes to another location, even if empty, washes out the neural memory of the previously seen offer in OFC: only the value of the offer that falls, or immediately fell, in the fovea is strongly encoded. This might imply that OFC does not hold a memory of all previously considered offers –just the one of the most recently fixated. These results are consistent with recent research showing that OFC only encodes the value of fixated offers, while other areas, such as the anterior cingulate cortex (ACC), might have a more detailed map of the relevant offers in the visual field and possibly perform a comparison (Butler et al., 2021). Therefore, these and our results suggest that OFC is unique in holding and evaluating the most recent offer and reactivating it when necessary for further evaluation. This view contrasts with the simultaneous encoding of offers previously reported in OFC in tasks where gaze remained fixed on a central fixation point (Ballesta & Padoa-Schioppa, 2019). An important difference of this study with ours is the lack of eye movements: as stimuli are passively projected onto the fovea, the task of maintaining previously encoded information might be simplified, thus making possible multiple offer’s value encoding. In contrast, in the more naturalistic scenario where gaze is required to foveate offers, it is possible that an association is made between offer and its location, such that later reactivation of a formerly presented offer requires, or is facilitated by, look-at-nothing gazing. Our results are also more in accord with other work showing that fluctuations of covert attention are translated into alternations between encoding one of two presented offers as a function of time, surprisingly independent of gaze (Rich & Wallis, 2016). Recent work where attention is decoupled from stimulus saliency shows that value encoding of well-known offers in single-neuron OFC is independent of attentional shifts (Zhang et al., 2022). It is possible that in this study overtraining makes simpler the encoding of value and thus gaze and attention can be decoupled. In contrast, in our case offers are more complex in number and variety but also in the requirement of having to combine two features (color for reward size and height for probability) to form an estimate of their value, and then it is possible that a more robust, gaze-centered working memory mechanism needs to be at place.

On the face of it, there is no clear reason why the brain would benefit from shifting gaze back to the previous location to reactivate a previously encoded offer. One possibility is that coupling between gaze and evaluation leverages brain machinery that allows for an efficient spatial representation of visual offers to be used for storage and comparison (Backen et al., 2018). Indeed, behavioral studies have proposed the gazing allows for memory index in spatial coordinates (Ferreira et al., 2008; Renkewitz & Jahn, 2012; Richardson & Spivey, 2000), therefore facilitating memory retrieval and decision making. Alternatively, it is also possible that economic decision making is necessarily embodied by linking gaze with cognitive processing, and that spontaneous associations between gazed spatial locations and offers form on the fly with no clear advantage beyond creating biases.

All in all, our results generally support the hypothesis of sequential value encoding (Hayden & Moreno-Bote, 2018), whereby specific brain allocates the neural resources to a single offer being scrutinized at a time. Recent theoretical work shows focused evaluation of offers is generally a much better strategy than dividing neural resources into several, or many, offers to be processed in parallel (Mastrogiuseppe & Moreno-Bote, 2022; Moreno-Bote et al., 2020). The one-at-a-time processing of offers during look-at-something and the specific re-activation of one offer at a time during look-at-nothing thus suggests a syntax of syllable-like neural operations that need to be executed sequentially in decision making.

## Methods

Two male rhesus macaques (*Macaca mulatta*) were trained to perform a two-alternative risky choice task (Strait et al., 2014). All procedures were approved by the University Committee on Animal Resources at the University of Rochester or the University of Minnesota and were designed and conducted in compliance with the Guide for the Care and Use of Animals of the Public Health Service. Recordings covered brain Areas 11 and 13, in orbitofrontal cortex (Fig. 3A). Data include 5971 trials correctly performed (2643 from subject 1, 3328 from subject 2) and 248 neurons (163 from subject 1, 85 from subject 2; the average number of simultaneously recorded cells per session is 40.75 ± 16.9 (mean ± s.d.) for subject 1, and 21.25±9.98 for subject 2), detailed number of trials and cells for each session and for each subject are reported in Supplementary Tables ST1 and ST2.

### 1. Behavioral task

The behavioral task consisted in the sequential presentation of two visual stimuli on the two opposite sides of the screen. Each visual stimulus indicated the size of a reward that could be chosen at the end of the trial by saccade to stimuli location (Fig. 1.A). The task starts with a first offer presentation (*offer 1*) displayed on the screen for 400 ms, followed by a first black screen delay time with duration 600 ms (*delay 1*); the same timings are used for a subsequent, second offer presentation (*offer 2*), respectively followed by a second, black screen delay time (*delay 2*). After the presentation of the two alternative offers, subjects were instructed to reacquire fixation (*re-fixate*), for at least 100 ms, at the center of the screen through the appearance of a central fixation dot. Following fixation, the choice could be indicated by the animal after the *choice-go* cue, which consisted of the simultaneous presentation of both offer stimuli in the same locations where they were previously shown. Choice was reported by shifting gaze to the preferred offer location and holding fixation on it for 200 ms (*choice-hold*). Fixation breaks during choice-hold periods led to a return to the choice report stage, giving subjects the chance to change their mind, thus to re-start fixation to either offer location for a duration of 200 ms. In trials with a valid choice report, the trial follows with the gamble outcome resolution: reward delivery for a successful outcome and lack of thereof for an unsuccessful outcome (see below for the probabilities of these events and the stimuli cues). Subjects were left free to direct their gaze during all task epochs except when instructed to fixate. Trials that took more than 7 seconds were considered inattentive and were not included in the analyses. Successfully rewarded trials were followed by visual *feedback* made of a white circle centered on the chosen offer stimulus.

Visual stimuli were presented on a 24” monitor with resolution 1024 pixels x 768 pixels, physical width 48.8 cm and height 36.6 cm at 57 cm from the eyes of the animals. The visual offers consisted of two vertical rectangles 300 pixels tall and 80 pixels wide (14.3 cm height and 3.8 cm width). Their centers were displaced right or left from the center of the screen by 256 pixels (12.2 cm,1/4 of total screen width). The presentation sites were randomized so that the first offer could be presented with equal probability to either side of the screen, and the second on the opposite side (the final count was that in 50.01% of the trials the first offer was presented on the left side of the screen). The colors of the rectangles indicated the magnitude 𝑚 and probability 𝑝 of the offered liquid reward. The magnitude of the reward was indicated by the color of the bottom part of the rectangle (gray: small, 125μL; blue: medium,165μL; green: large, 240μL). In all analyses, reward magnitudes 𝑚 are reported and used in nominal units (1 = small, 2 = medium, 3 = large reward). The height of the bottom fraction of the stimuli indicated the (success) probability 𝑝 of obtaining the liquid reward (with magnitude 𝑚 as indicated by its color) if the offer was chosen. Whenever 𝑝 was <1, the stimuli top fraction was colored in red, with its height indicating the complementary probability 1 − 𝑝 for unsuccessful outcome. We define the expected value of an offer as the product of its probability times its magnitude: 𝐸𝑉 = 𝑚𝑝. To consider the trial-to-trial variability of offer value, we also defined the variance 𝜎^2^ = 𝑚𝑝(1 − 𝑝), also referred to as offer risk. Offers having small rewards were always sure, 𝑝 = 1 (safe option). Reward probabilities for the medium and large magnitude offers were randomly and independently drawn from a uniform distribution in the unit interval, whose presentation was limited in size only to the resolution of screen pixel size. Reward magnitudes (small, medium, or large) of the first and second offer were randomized across trials so that 1/8 were safe, and for the remainder 7/8, offers could be either medium or large in equal fractions (yielding a total of: 12.15% small, safe offers; 43.93% medium sized gamble offers, 43.92% large sized gamble offers). Trials were interleaved with an 800ms idle inter-trial interval when subjects were not given instructions and the screen was left blank.

Eye position was sampled at 1,000 Hz by an infrared eye-monitoring camera system (SR Research). Stimuli were controlled by a computer running Matlab (Mathworks) with Psychtoolbox (Brainard, 1997) and Eyelink Toolbox (Cornelissen et al., 2002).

### 2. Behavioral data analyses

In all analyses, we pool trials where the first offer was on the left with trials where the first offer was on the right by mirroring in the latter behavioral variables and eye tracking data along the vertical axis. Therefore, in all analyses and figures the first offer is anchored on the left presentation site, since we did not find appreciable difference between trials with opposite order of presentation (Supp. Fig. S1B). For time resolved analyses, time epochs are aligned to task-related event times (e.g., *offer 1* time runs from stimulus onset up to the end of its presentation). For time epochs with random duration (e.g., for *delay 2*, due to trial-to-trial variation in fixation re-acquisition time), we limited the end of task-relevant time epochs to the duration of shortest trial across all data (in both monkeys and sessions), so that all analyzed data points were covered by all trials.

#### 2.1 Behavioral performance

We quantify the behavioral performance on the value-based decision-making task by regressing the fraction of choices for Right offer side (𝑐ℎ𝑅 = 1 if chosen offer is on Right side; 𝑐ℎ𝑅 = 0 if chosen offer is on the Left side) by the difference in the expected value of the two offers 𝐸𝑉_𝑅_ − 𝐸𝑉_𝐿_ (Fig. 1B). We fit choice data to the model: logit(𝑐ℎ𝑅) = 𝛽_0_ + 𝛽_1_(𝐸𝑉_𝑅_ − 𝐸𝑉_𝐿_), and assess the significance of linear interaction term 𝛽_1_ via *F*-Statistic test to compare the model fit vs a constant mean model (with 𝛽_1_ = 0), ***𝑝<0.001.

#### 2.2 Analyses of eye position and gaze shifts

The spatial distribution of horizontal eye position across task epochs (Fig. 1C, top) is computed by counting occurrences across trials with spatial resolution 0.14 cm at a temporal resolution of 1 ms. The counting is then applied by splitting trials where subjects mainly looked either left or right screen side (Fig. 1C, bottom). We call a trial *LookL* or *LookR* based on whether the average horizontal eye position in 10 ms bins is negative or positive, respectively. The 2D spatial distribution of eye position on the screen is reported as heatmaps around the center of the visual scene (Fig. 1.D), binned with resolution 0.14 cm, smoothed with a gaussian filter (𝜎_𝑥_ = 𝜎_𝑦_ = 5 bins, 0.7 cm). The 2D analysis is initially run for each task epoch and by pooling data across all sessions from the two subjects (Fig 1D), then repeated by separating trials by best offer 𝐸𝑉 (best L: 𝐸𝑉_𝐿_> 𝐸𝑉_𝑅_, or best R: 𝐸𝑉_𝑅_> 𝐸𝑉_𝐿_; Fig. 1E), by separating data from the two subjects (Supp. Fig. S2A), for the two orders of presentation (first = L, or first = R; Supp. Fig. S2B), and for different choices (choice = L, or choice = R; Supp. Fig. S2C). Results for different conditions are subtracted to highlight differences between best offer cases (best R − best L, Supp. Fig. S2D) and chosen offers (choice R − choice L, Supp. Fig. S2E).

We analyzed horizontal gaze shifts, which we defined as monotonic variations in the horizontal eye position with duration of at least 25ms (Supp. Fig. S1C). The shifts are labeled by horizontal direction (left/right) based on the sign of their first order discrete-time derivative (left if the eye position derivative is negative), at 1ms resolution. We show the time histograms of horizontal gaze shifts both for shifts within each visual hemifield and by also including midline-crossing (Supp. Fig. S1C).

#### 2.3 Probability of choice as a function of offer expected values and looking times

For each task-related time epoch (*offer 1*, *delay 1*, etc.) and for each trial, we computed the fraction of time looking at the right screen side, 𝑓_𝑅_ = 𝑡_𝑅_/(𝑡_𝑅_ + 𝑡_𝐿_), where 𝑡_𝑅_ is the total time that the subject spends looking at the right screen side, and 𝑡_𝐿_ is the total time looking at left screen side in that epoch and trial.

We further analyzed the choice variable *chR* (set to 1 if choice is right, 0 if left) as a function of the difference in expected value of the two offers 𝐸𝑉_𝑅_ − 𝐸𝑉_𝐿_ (Fig. 2A). The choice variable was fit using least squares method to the model logit(𝑐ℎ𝑅) = 𝛽_0_ + 𝛽_1_(𝐸𝑉_𝑅_ − 𝐸𝑉_𝐿_), with a resolution of 0.2 nominal reward units. First, we run the analysis including all trials, then we split trial pools where subject mainly look left (𝑓_𝑅_ < 0.5; light red in Fig. 2A), or right (𝑓_𝑅_ > 0.5; dark red in Fig. 2A). The ±95% C.I. is computed via the estimation of inverse cumulative t-distribution given the coefficients (𝛽_0_, 𝛽_1_), their covariance matrix and standard errors of the fit. The differences in slope show that the eye position shifts the logistic relationship between choice and 𝐸𝑉 difference, hence that eye position affects the choice beyond the 𝐸𝑉 differences.

Following a similar logic, we predict the chosen offer 𝑐ℎ𝑅 across trials using the logistic regression model logit(𝑐ℎ𝑅) = 𝛽_0_ + 𝛽_1_ (𝑡_𝑅_ − 𝑡_𝐿_), i.e. by including as regressor the difference in right vs left screen looking times 𝑡_𝑅_ − 𝑡_𝐿_ within each trial (Fig. 2B). The difference in looking times 𝑡_𝑅_ − 𝑡_𝐿_ is normalized to the maximum of its absolute value across trials (maximum is taken across data in all sessions and in both subjects), and therefore the value of this regressor always falls within [-1, +1]. The ±95% C.I. is computed via estimation of inverse cumulative t-distribution as before. We find a significant relationship between choice and difference in inspection time for the two screen sides in all task epochs. The same analysis is repeated for trial pools where first offer is best (𝐸𝑉_𝐿_ > 𝐸𝑉_𝑅_, light purple in Fig. 2B) and second offer is best (𝐸𝑉_𝑅_ > 𝐸𝑉_𝐿_, dark purple in Fig. 2.B). We found a shift in the relationship between choice and difference in looking times, indicating that choice for either offer was more likely if the same offer was looked at for longer time, and if it had best 𝐸𝑉.

Further than this, we refine the above models by combining task variables in the logistic model 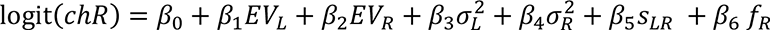 (Fig. 2C), with the following trial-by-trial regressors: expected value of left and right offers 𝐸𝑉_𝐿_ and 𝐸𝑉_𝑅_ (computed as 𝐸𝑉 = 𝑚𝑝); risk, or variance of left/right offers, 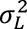 and 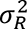 (computed as 𝜎^2^ = 𝑚𝑝 (1 − 𝑝)); order of offers presentation 𝑠_𝐿𝑅_ (+1 if first offer is left, −1 first offer is right); and fraction of time looking at right screen side 𝑓_𝑅_ = 𝑡_𝑅_/(𝑡_𝑅_ + 𝑡_𝐿_). To compare the magnitudes of the regressors, we normalized 𝐸𝑉 and 𝜎^2^to their maximum value across all trials all sessions, and in both subjects, so for the analyses on the whole data set their trial-by-trial values ranged within unitary values (in the set [0, 1]). Results for the regression of EV and 𝜎 only are reported in Supplementary Table ST3. Results are also shown separately for the two subjects (Supp. Fig. S4A). To test the effect of the fraction of looking time (𝑓_𝑅_) on the choice (*chR*), we added to the previous logistic regression model the fractions of looking at the right side in each of the following relevant task epochs: (*offer 1*, *delay 1*, *offer 2*, *delay 2* (Fig. 2D), showing that 𝑓_𝑅_ in *delay 2* has significant impact on the choice (**𝑝 < 0.01) even when other regressors are included. We performed an addition analysis by redefining the looking times 𝑡_𝑅_ and 𝑡_𝐿_ as the total amount of time when the gaze of the subjects is inside the physical locations of the offers (or when they are left empty) (Supp. Fig. S3B). The locations of the visual stimuli are: left side, horizontal coordinates −7.5±1.5cm; right side, horizontal coordinates +7.5±1.5cm; Left and right sides: vertical coordinates between ± 6 cm. This allowed us to determine whether gaze spatial precision is important for the encoding of the stimulus (gazing to the offer vs looking to the screen side where the offer was located). We find no qualitative difference in our results when using screen vs offer looking times (Supp. Fig. S3B; compare with Fig. 2C).

### 3. Neuronal data recordings

Animals were habituated to laboratory conditions and then trained to perform oculomotor tasks for liquid reward. We approached Areas 11 and 13 through a standard Cilux recording chamber (Crist Instruments). A small prothesis for holding the head was used. Position was verified by magnetic resonance imaging by aligning white and gray matter scans to standard anatomical atlas with the aid of a Brainsight system (Rogue Research Inc.). Neuroimaging was performed using a 3T MAGNETOM Trio Tim using 0.5 mm wide voxels. Animals received appropriate analgesics and antibiotics after all procedures. Throughout both behavioral and physiological recording sessions, the chamber was kept sterile with regular antibiotic washes and sealed with sterile caps. Single contact electrodes (Frederick Haer & Co., impedance range 0.8 to 4 MU) were lowered using a microdrive (NAN Instruments) until waveforms of between one and three neuron(s) were isolated. Individual action potentials were isolated on a Plexon system (Plexon, Inc.). Following a settling period, all active cells were recorded. Cells were sorted offline manually by trained electrophysiologists; no automated sorting tools were used. Neurons were selected for study solely based on the quality of isolation; they were never preselected based on task-related response properties. All collected neurons for which we managed to obtain at least 300 trials were analyzed; no neurons that surpassed our isolation criteria were excluded from analysis.

### 4. Analyses of neural data

Following the manual isolation of units, spiking activity data were preprocessed by thresholding, quantization, and subsampling to 1kHz resolution. The resulting data have an average spike rate of 1.46 ± 2.38 (mean ± s.d.) spikes/s. Given the sparsity of the data, to improve the detectability of encoding of task variables, we always applied the analysis of neural data by using a time-resolved strategy based on boxcar sliding window.

#### 4.1 Neural spiking and the encoding of reward expected value

For each of the recorded units, we computed the time-averaged spike count 𝜂 in time windows of 200ms, in every 10 ms bins (consecutive time windows have 95% overlap; time windows start at every 10 ms bin). These settings were chosen because they provided a good trade-off between the temporal resolution of our analysis and large enough time windows to compute spike rates given the sparsity of OFC activity. In addition, these settings were in line with ranges used in previous studies of OFC activity (Conen, Padoa-Schioppa, 2015; Maisson, D. et al., 2021). The spike rate in each time window, for each neuron, and in each 10 ms bin was fit to the linear model 𝜂 = 𝛽_0_ + 𝛽_1_𝑥, where 𝑥 was either the 𝐸𝑉 of the left offer, 𝐸𝑉_𝐿_, or right offer, 𝐸𝑉_𝑅_. This time-resolved, cell-by-cell analysis was performed using three different sets of trials: (1) using all available trials, i.e. neglecting where the animal is looking; (2) using all trials within a session where the animal mainly looks at the left side of the screen in a given bin (defined as trials where the average gaze position within the 10 ms time bin is negative); ; and (3) trials where the animal mainly looked at the right side (average gaze position is positive). The sets of trials (2) and (3) allowed to study the modulatory effect of gaze on the encoding of the value of the offers by distinguishing trials where subjects mainly *‘Look Left = LookL’* from trials where subjects mainly *‘Look Right = LookR’,* respectively (the bottom panel of Fig. 1C shows the fraction of trials for *LookL / LookR* for each session, in each time bin). The number of available trials could vary across sessions (𝑛 = 746.38 ± 87.29 mean ± s.e.m trials per session, 𝑛 = 5971 in total; Supp. Table ST1 for exact numbers for each session) but coincided for simultaneously recorded cells. Given that both subjects shared the look-at-something and look-at-nothing gaze patterns for the two screen sides, we considered looking conditions as the opposite sides of the screen (*LookL / LookR*), which also allowed to maximize the usage of available trials. We did not consider looking conditions as based on perfect alignment of gaze position over stimuli presentation sites as the subjects gaze did lack precision in reaching exact stimuli locations (e.g. *delay 2* in Fig. 1D,E; Supp. Fig. S2). Such looking conditions drastically reduce the number of available trials, preventing us from applying neural encoding analyses, especially at delay times (at *delay 1* time the fraction of available trials is 9.79% of the total for *LookL*, 1.75% for *LookR*; at *delay 2* time it is 3.76% for *LookL*, 7.43% for *LookR*; see Supp. Table ST2 for available trials at opposite screen sides).

To compute the fraction of significant cells, we pooled cells across all sessions for the two subjects (𝑛 = 31 ± 5.85 mean ± s.e.m. cells per session, 𝑛 = 248 in total; Supp. Table ST1 for exact numbers per session). The significance of each cell in encoding the offer 𝐸𝑉 in each time bin is assessed via *F-*Statistic tests comparing the linear model vs constant model, thus assessing whether the linear coefficient 𝛽_1_ is significantly different from zero (𝑝 < 0.05). From this, we estimated the fraction of significant cells over the total number of available cells (for both monkeys, Fig. 3B; for the two monkeys separately, Supp. Fig. S5). In our results we do not find qualitative change if shorter (150ms) or longer (250ms) time windows are used for spike rates (Supp. Fig. S6), nor if we applied baseline normalization (for each cell, for each trial, we subtracted the spike rate in each 10ms bin by average spike rate in *pre-offer 1*, Supp. Fig. S7).

Although we mostly showed results for the neural encoding in terms of fraction of significantly encoding cells, we also show the coefficient of determination 𝑅^2^ for all the regressions that we implemented (Supp. Fig. S8A). In addition, we repeated our significant tests based on *F*-tests by(non-parametric) permutation tests on the values of 𝑅^2^(Supp. Fig. S8B). In the permutation test, the trial order of 𝐸𝑉s and spike-counts are shuffled independently 𝑛 = 1000 times in each 10 ms time bin, for each cell, in each session, for the two subjects. Then, the empirical R^2^ (based on the not shuffled data) is compared to the 𝑛 = 1000 shuffled 𝑅^2^ values, and the p-value is defined as the fraction of shuffles with 𝑅^2^ larger than empirical 𝑅^2^(Nogueira et al., 2017). Finally, we counted the cells with significant encoding (𝑝 < 0.05). The results for the permutation test (Supp. Fig. S8B) qualitatively match results run via *F*-statistic tests (Fig. 3B,D).

To provide our bin-resolved analysis with statistical robustness, we did not naively consider significance of the fraction of cells for single bins, as many of those bins could be significant just by chance (i.e., the eventuality that the fraction of significant cells in any time bins crossed the 95^th^ percentile significance threshold due to spurious fluctuations). To address this issue, we performed a cluster-based run-length analysis as follows (Butler et al., 2021). Here, we defined the length of a run as the number of consecutive significant time bins (significant here means that in the current time bin we have a fraction of significant cells exceeding the 95^th^ percentile of the fraction of cells from shuffled data). We considered that a run length was significantly longer than expected by chance if its length exceeded the 95^th^ percentile of run lengths obtained from shuffled data. The time bins involved in significantly long runs are shown by the solid line at the bottom of each panel in Fig. 3B-D, in Fig.4, and in Supp. Figs. S4-S10.

In all our neural data analyses we compared with the respective control results obtained over trial-order shuffled data. For each time bin, we shuffled the trial order of spike rates and expected values independently for each cell, to destroy the correlation between spike rates, expected values and eye position. For each of the 𝑛 = 1000 independent shuffles generated, we computed the fraction of cells that encoded either the left or right expected values. The 1000 fractions were used to build the null-hypothesis distribution. Finally, the 95^th^ percentile (right tail) of the null-hypothesis distribution is used to test at 5% significance level if the observed fraction of significant cells is significantly larger than the one expected by chance. We chose a right-tailed test because we consider only meaningful fractions of significant cells larger than expected by chance, not smaller. For illustration purposes, we display the 5^th^-to-95^th^ percentile of the null-hypothesis distribution as shaded areas in Fig. 3B-D, in Fig.4 and in Supp. Figs. S4-S10.

We repeated the analysis of Fig. 3D by controlling for the unbalanced number of trials for the conditions *‘LookL’* and *‘LookR’* (see Fig. 1C, bottom). To implement this, we performed a sub-sampling trial method such that the number of trials for each condition is the same, before performing statistical comparisons (Supp. Fig. S9). In each time bin, the number of trials for both *LookL* and *LookR* is set to 𝑛(𝑡) = min (number of trials *LookL*, number of trials *LookR*) in each session (e.g. 𝑛(𝑡) = min(60, 535)). The number of subsets at each time bin is given by 𝑚(𝑡) = ⌈𝑁/𝑛(𝑡)⌉, with *N* the total number of trials in each session (e.g., 𝑚(𝑡) = ⌈595/60⌉ = 10). Then, the average fraction of significant cells is computed for each subset, and finally the fractions for all 𝑚(𝑡) subsets are averaged. We show that the main results (Fig. 3D) are found by sub-sampling (Supp. Fig. S9C), yielding equivalent overall levels of 𝑅^2^ for *LookL* and *LookR* (Supp. Fig. S9D).

### 4.2 Difference in encoding 𝑬𝑽_𝑳_ versus 𝑬𝑽_𝑹_ as a function of time, and difference in encoding 𝑬𝑽_𝑳_ or *𝑬𝑽_𝑹_* when looking at different looking sides: LookL versus LookR

We tested whether in any task-related epoch (e.g., *offer 1*, *delay 1*, etc.), cells showed more encoding of 𝐸𝑉_𝐿_ or 𝐸𝑉_𝑅_. We first consider all the trials, regardless of where the monkeys directed their gaze (below, we conditioned the same analysis on the gaze of the subjects). For each task-related epoch, we first built two 𝑛_𝑐𝑒𝑙𝑙𝑠_ × 𝑛_𝑡𝑖𝑚𝑒𝑏𝑖𝑛𝑠_ binary matrices, 𝑀_𝐿_ and 𝑀_𝑅_. In each matrix, the element 𝑚_𝑖,𝑗_ = 1 if cell 𝑖 at time bin 𝑗 has a significant encoding of the left or right 𝐸𝑉 (𝑝 < 0.05, defined by F-tests on the linear term interaction coefficient described in previous Section 4.1), 𝑚_𝑖,𝑗_ = 0 otherwise.

We compute the fraction of time bins with significant encoding of either expected value by averaging the matrices 𝑀_𝐿_ and 𝑀_𝑅_ along time dimension in each task epoch, yielding vectors 𝑢_𝐿_ and 𝑢_𝑅_ with size 𝑛_𝑐𝑒𝑙𝑙𝑠_ × 1. In Fig. 3C we report average ± s.e.m. across cells of the vectors 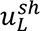 and 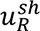. For each of the two cases, we computed the 95^th^ percentile values from the distribution of the fraction of bins averaged across cells 𝑢_L_^𝑠ℎ^ and 𝑢_R_^𝑠ℎ^ for 𝑛 = 1000 shuffles of the trials order, to be used as significance threshold to be exceeded (in Fig. 3C, non-significant bars are reported in white, with colored frame). The two vectors 𝑢_𝐿_ and 𝑢_𝑅_ were compared via non-parametric tests (Wilcoxon signed rank test; *p<0.05, **p<0.01 and ***p<0.001 in Fig. 3C).

After this, we analyzed the fraction of neurons that encoded the left and right expected values depending on the subject’s gaze in each time bin within a task-related epoch. We divided trials into two groups: *LookL* trials, where average gaze position is < 0 in the 10 ms time bin in each of those trials; and *LookR* trials, where average > 0. Focusing on either offer 𝐸𝑉_𝐿_ and 𝐸𝑉_𝑅_, we re-run the neural encoding analysis again to compute the fraction of cells significantly encoding for either looking side (𝐸𝑉_𝐿_ *LookL* and *LookR*, Fig. 3D, top; 𝐸𝑉_𝑅_ for *LookL* and *LookR*, Fig. 3D, bottom). For both the 𝐸𝑉𝑠 and for the two trial pools, we computed again the vectors 𝑢_𝐿_ and 𝑢_𝑅_ as above, called them 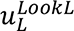 and 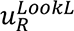 for the *LookL* trials, and 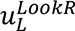 and 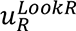 for the and *LookR* trials. Like in Fig. 3C, we report average ± s.e.m. across cells of the vectors 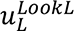, 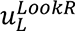 (Fig. 3E, top), and 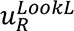, 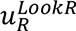 (Fig. 3E, bottom) (non-significant bars are reported in white, with colored frame). We compared the encoding of left expected value 𝐸𝑉_𝐿_ when the animal mostly looked at left versus the encoding when it mostly looked at right, that is, we compared 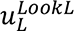 vs 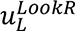, again, via non-parametric tests (one-sided Wilcoxon signed rank test; *p<0.05, **p<0.01 and ***p<0.001 in Fig. 3E, top). This time we use one-sided tests as we are interested in *LookL* significance being more prominent than in *LookR*, and not just significantly different in their mean value. The same is done for the effect of gaze on the encoding of right expected value 𝐸𝑉_𝑅_, i.e., we applied non-parametric tests to compare 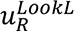 vs 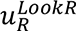 (Fig. 3E, bottom), this time assessing whether *LookR* was larger than *LookL*, as we wanted to test stronger encoding of 𝐸𝑉_𝑅_.

### 4.3 Neural encoding at delay times conditioned on previous looking side

To determine whether memory traces and reactivation of activity in OFC are mostly due to trials where animals did not change their gaze (e.g., whether the animals look at the left side during *delay 2* because it did not look at the right side during *offer 2*), we repeated the encoding analysis during the delay periods but making sure that we used only trials where the animal actually looked at the offer immediately before them. We first focused on *delay 1* time and compared *LookL* vs *LookR* by conditioning on trials when subjects were previously looking at the left during the last 200 ms of *offer 1* (looking at the left is of course the most common gazing pattern). We find that the subject *LookL* during the last 200 ms of *offer 1* on 80.21% of the total amount of trials, they keep *LookL* during the last 200 ms of *delay 1* on 47.95% of total amount of trials, while they switch to *LookR* during the last 200 ms of *delay 1* on 32.26% of the total trials. For the two cases we applied independently the regression of 𝐸𝑉_𝐿_ and 𝐸𝑉_𝑅_, as described in previous Section 4.2 (Fig. 3F). We find that the encoding of the first offer 𝐸𝑉_𝐿_ persisted if the subjects keep looking at the left side of the screen during *delay 1*. Next, we focused our analyses on *delay 2* time and we compared looking conditions for trials when subjects were looking to the right during the last 200 ms of *offer 2*, which is the most common pattern (78.8% of the total amount of trials). Again, we applied the time-resolved regression of offer 𝐸𝑉s to compare trials when subjects shifted to *LookL* (36.48% of total amount of trials) versus the trials when subjects kept *LookR* (42.32% of total amount of trials) during *delay 2* (Fig. 3G). We strikingly find significant increase in the encoding of left offer 𝐸𝑉_𝐿_ when subjects look back to first offer location during *delay 2*, despite the screen was left blank; and this reactivation occurred even when the second offer was looked at, which means that reactivation is robust to distractors. For 𝐸𝑉_𝑅_, just like for 𝐸𝑉_𝐿_ in *delay 1*, we find that its encoding persists only if the subjects keep staring at ipsilateral screen location. The analyses were run also for less common, complementary cases (*LookR* during *offer1* and *LookL* during *offer 2*), shown in Supp. Fig. S4A, Supp. Fig S4B.

As a further analysis, we computed the correlation of regression weights for the linear regression weights 𝛽_1_ for the spike rate and 𝐸𝑉 of the offers (𝜂 = 𝛽_0_ + 𝛽_1_𝐸𝑉). We compared the weights for the two look-at-nothing cases (*LookL* / *LookR* during *delay 2*) and the respective look-at-something cases (*LookL* / *LookR* during *offer 1* / *offer 2*). In this analysis we computed the regression weights in the last 200 ms of each epoch time (*offer 1* / *offer 2* / *delay 2*) by considering the average spike rate in the whole windows (instead of using a time-resolved approach used in the remainder of this section). This choice is motivated by the observation that both eye position and neural encoding results show steady patterns in these time windows. Lastly, we computed Spearman correlations between the 𝛽_1_ of each cell in different pairs of task scenarios, i.e., for ipsilateral looking cases (Supp. Fig. S11A) and contralateral cases (Supp. Fig. S11B).

### 4.4 Neural encoding of offer subjective value and the choice

We further tested a more complete model where we applied the regression of both offer Subjective Values (𝑆𝑉) and the choice variable 𝑐ℎ𝑅 (𝑐ℎ𝑅 = 1 if choice is for right offer, 0 otherwise). The *SV*s are computed by using behavioral data in each session, as 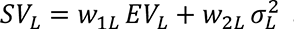 and 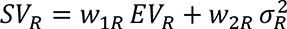, regressing the choice variable as 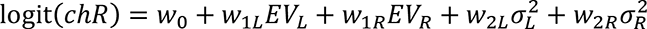. For the analysis of neural data, we fit the linear model 𝜂 = 𝛽_0_ + 𝛽_1_𝑆𝑉_𝐿_ + 𝛽_2_𝑆𝑉_𝑅_ + 𝛽_3_𝑐ℎ𝑅 to spike rate 𝜂 in each 10 ms bins, using a sliding time window of 200 ms. In this analysis we followed the same methodology as for the time-resolved analysis previously run independently for the encoding of 𝐸𝑉_𝐿_ and 𝐸𝑉_𝑅_ described in Section 4.1. To investigate the modulatory effects of gaze position, we repeated the same analysis by either including all trials (Fig. 4A, top), or focusing on *LookL* trials (Fig. 4A, middle), and on *LookR* trials (Fig. 4A, bottom). Interestingly, we found that our main results for the two 𝑆𝑉s qualitatively matched results for the encoding of the two 𝐸𝑉s in isolation, even when including the choice variable, which in turn is significantly encoded in the late task epochs. We first assess the mean fraction of significant cells by comparing them with the 95^th^ percentile of the same quantities computed over 𝑛 = 1000 shuffles of the trial order, to be used as significance threshold (in Figs. 4B and 4D, non-significant bars are reported in white, with colored frame). Then, for the encoding of 𝑆𝑉s, we compared the fraction of significant bins in *LookL* and *LookR* (Fig. 4D, top and middle) via one-sided Wilcoxon signed-rank tests, to test that either 𝑆𝑉 had higher encoding for ipsilateral inspection (following the derivation of the vectors 𝑢_𝐿_ and 𝑢_𝑅_ described in Section 4.2). For the encoding of 𝑐ℎ𝑅 (Fig. 4D, bottom), we compared *LookL* and *LookR* (again using 𝑢_𝐿_ and 𝑢_𝑅_ computed as in Section 4.2) via two-tailed Wilcoxon signed-rank tests to test that the two conditions showed non-zero mean difference.

## Code availability

Code and data is available at the DOI: <u>10.12751/g-node.72yf9s</u>.

## Acknowledgements

We appreciate Ramon Nogueira for discussion about core methods.

This project was supported by grants from the Howard Hughes Medical Institute (HHMI; Ref: 55008742), The Bial Foundation (Ref: 106/2022), Ministerio de Ciencia e Innovación (Ref: PID2020-114196GB-I00/AEI) and ICREA Academia (2022) to R.M.B.

## Author contributions

D.F., R.M.B. conceptualized and designed the analyses. B.H. ideated the task. D.F. analyzed the data. B.H., T.C.P., M.Z.W. collected the data. D.F., B.H., R.M.B. wrote the manuscript.

