## Supplementary Materials for "Gaze-centered gating and reactivation of value encoding in orbitofrontal cortex"

### Supplementary Information

| Subject 1 | | | | Subject 2 | | | |
|---|---|---|---|---|---|---|---|
| **Area** | **Session** | **#cells** | **# trials** | **Area** | **Session** | **#cells** | **#trials** |
| 13 | 1 | 51 | 643 | 11 | 1 | 18 | 1015 |
| 13 | 2 | 59 | 700 | 11 | 2 | 32 | 323 |
| 11 | 3 | 24 | 697 | 11 | 3 | 9 | 1084 |
| 11 | 4 | 29 | 603 | 11 | 4 | 26 | 906 |
| **Total** | | 163 | 2643 | **Total** | | 85 | 3328 |
| **Mean±sem** | | 40.75±8.45 | 660.75±23.28 | **Mean±sem** | | 21.25±4.99 | 832.00±173.58 |
| **Subjects 1:2** | | **Total** | **#cells** | 248 | **#trials** | 5971 | |
| | | **Mean±sem** | **#cells** | 31±5.85 | **#trials** | 746.38±87.29 | |

**Table ST1. Number of sessions, cells, and trials for the two subjects.**

| | ***first=L*** | ***first=R*** | ***choice=L*** | ***choice=R*** | $EV_L>EV_R$ | $EV_L<EV_R$ | $EV_L=EV_R$ |
|---|---|---|---|---|---|---|---|
| **Subject 1** | 1310<br>(49.56%) | 1333<br>(50.44%) | 1174<br>(44.42%) | 1469<br>(55.58%) | 1298<br>(49.11%) | 1294<br>(48.96%) | 51<br>(1.93%) |
| **Subject 2** | 1676<br>(50.36%) | 1652<br>(49.64%) | 1790<br>(53.79%) | 1538<br>(46.21%) | 1660<br>(49.88%) | 1593<br>(47.87%) | 75<br>(2.25%) |
| **Total** | 2986<br>(50.00%) | 2985<br>(50.00%) | 2964<br>(49.64%) | 3007<br>(50.36%) | 2958<br>(49.54%) | 2887<br>(48.35%) | 126<br>(2.11%) |
| ***offer1 LookL***<br>***LookR*** | **78.83%**<br>21.17% | 18.43%<br>**81.57%** | **87.11%**<br>12.89% | **73.44%**<br>57.83% | **84.62%**<br>15.38% | **76.20%**<br>23.80% | **69.84%**<br>30.16% |
| ***delay1 LookL***<br>***LookR*** | 44.78%<br>55.22% | 35.51%<br>64.49% | **62.77%**<br>37.23% | 46.65%<br>**53.35%** | **60.38%**<br>39.62% | 49.07%<br>**50.93%** | 49.21%<br>50.79% |
| ***offer2 LookL***<br>***LookR*** | 18.18%<br>**81.82%** | **75.78%**<br>24.22% | 32.32%<br>**67.68%** | 10.31%<br>**89.69%** | 30.34%<br>**69.66%** | 12.05%<br>**87.95%** | 19.84%<br>**80.16%** |
| ***delay2 LookL***<br>***LookR*** | 43.30%<br>56.70% | 41.34%<br>58.66% | **67.17%**<br>32.83% | 35.11%<br>**64.89%** | **63.65%**<br>36.35% | 38.32%<br>**61.68%** | 48.41%<br>51.59% |
| ***choice LookL***<br>***LookR*** | 49.30%<br>50.70% | 50.25%<br>49.75% | **99.90%**<br>0.10% | 0.17%<br>**99.83%** | **75.08%**<br>24.92% | 23.56%<br>**76.44%** | 54.76%<br>45.24% |

| | | | | | |
|---|---|---|---|---|---|
| ***offer1 LookL*** | **80.21%** | ***delay1 LookL*** | 54.63% | ***(Fig. 3F) offer1 LookL and delay1 LookL***<br>***offer1 LookL and delay1 LookR*** | **47.95%**<br>32.26% |
| ***offer1 LookR*** | 19.79% | ***delay1 LookR*** | 45.37% | ***(Fig. S10A) offer1 LookR and delay1 LookL***<br>***offer1 LookR and delay1 LookR*** | 6.68%<br>13.11% |
| ***offer2 LookL*** | 21.20% | ***delay2 LookL*** | 50.98% | ***(Fig. S10B) offer2 LookL and delay2 LookL***<br>***offer2 LookL and delay2 LookR*** | 14.50%<br>6.70% |
| ***offer2 LookR*** | **78.80%** | ***delay2 LookR*** | 49.02% | ***(Fig. 3G) offer2 LookR and delay2 LookL***<br>***offer2 LookR and delay2 LookR*** | 36.48%<br>**42.32%** |

**Table ST2. Number of trials (% fractions) for different task conditions. Fraction of trials for the two different screen sides (*LookL*/*LookR*) in main task epochs.**

Trials with $EV_L$=$EV_R$ are cases where both offers were safe ($EV_L$=$EV_R$=1). To capture meaningful patterns, offer presentation times and delay times (*offer1/delay1/offer2/delay2*) are cut to the last 200 ms of their duration. For *choice* time we used the first 440 ms of *choice-hold* epoch time allowing to capture almost all choices (with rare, 0.27% occasional misses). Trials are sorted with reference to first offer anchored on the left screen side (trials with first offer on right screen side are mirrored).

| $EV_L, \sigma_L^2$ | **Subject1** | | | | **Subject2** | | | |
|---|---|---|---|---|---|---|---|---|
| $EV_R, \sigma_R^2$ | session1 | session2 | session3 | session4 | session1 | session2 | session3 | session4 |
| $w_0$ | 1.06*** | 0.24 | 0.76* | -0.03 | -0.21 | -0.11 | -0.15 | 0.31 |
| $w_{1L}$ | 1.52*** | 1.72*** | 1.88*** | 2.10*** | 1.84*** | 2.26*** | 2.93*** | 2.41*** |
| $w_{2L}$ | 4.14*** | 4.13*** | 5.33*** | 6.21*** | 5.32*** | 6.20*** | 6.59*** | 5.84*** |
| $w_{1R}$ | 1.36*** | 1.60*** | 1.51*** | 2.21*** | 2.16*** | 2.72*** | 3.08*** | 2.09*** |
| $w_{2R}$ | 2.56*** | 3.78*** | 4.72*** | 6.19*** | 5.44*** | 4.90*** | 6.93*** | 6.04*** |

**Table ST3. Weights for the logistic regression of choice and offer value, risk.** The regression is applied for left offer $(EV_L, \sigma_L^2)$ and right offer $(EV_R, \sigma_R^2)$ as $\text{logit}(chR) = w_0 - w_{1L}EV_L - w_{2L}\sigma_L^2 + w_{1R}EV_R + w_{2R}\sigma_R^2$. Note: $EV$ range is $[0, 3]$; $\sigma^2$ range is $[0, 0.75\ (= 0.5 \cdot 0.5 \cdot 3)]$. Significance: *p<0.05, ***p<0.001.

| | ***offer1*** | ***delay1*** | ***offer2*** | ***delay2*** | ***re-fixate*** | ***choice-go*** | ***ch-hold*** |
|---|---|---|---|---|---|---|---|
| all trials | $\beta_0$ 0.04<br>$\beta_1$ 1.74$^{***}$ | $\beta_0$ 0.04<br>$\beta_1$ 1.74$^{***}$ | $\beta_0$ 0.04<br>$\beta_1$ 1.74$^{***}$ | $\beta_0$ 0.04<br>$\beta_1$ 1.74$^{***}$ | $\beta_0$ 0.04<br>$\beta_1$ 1.74$^{***}$ | $\beta_0$ 0.04<br>$\beta_1$ 1.74$^{***}$ | $\beta_0$ 0.04<br>$\beta_1$ 1.74$^{***}$ |
| $f_R > 0.5$ | $\beta_0$ 0.43<br>$\beta_1$ 1.55$^{***}$ | $\beta_0$ 0.34<br>$\beta_1$ 1.66$^{***}$ | $\beta_0$ 0.28<br>$\beta_1$ 1.67$^{***}$ | $\beta_0$ 0.41<br>$\beta_1$ 1.69$^{***}$ | $\beta_0$ 0.32<br>$\beta_1$ 1.76$^{***}$ | $\beta_0$ 0.30<br>$\beta_1$ 1.75$^{***}$ | $\beta_0$ 7.53<br>$\beta_1$ 2.76$^{**}$ |
| $f_R < 0.5$ | $\beta_0$ -0.11<br>$\beta_1$ 1.80$^{***}$ | $\beta_0$ -0.13<br>$\beta_1$ 1.75$^{***}$ | $\beta_0$ -0.59<br>$\beta_1$ 1.70$^{***}$ | $\beta_0$ -0.51<br>$\beta_1$ 1.60$^{***}$ | $\beta_0$ -0.18<br>$\beta_1$ 1.69$^{***}$ | $\beta_0$ -0.17<br>$\beta_1$ 1.69$^{***}$ | $\beta_0$ -5.92<br>$\beta_1$ 0.72 |

**Table ST4. Weights for the logistic regression of choice and difference in EV of the two offers.** Regression weights ($\beta_1$) and significance (**p<0.01, ***p<0.001) for results in Fig. 2A.

| | ***offer1*** | ***delay1*** | ***offer2*** | ***delay2*** | ***re-fixate*** | ***choice-go*** | ***ch-hold*** |
|---|---|---|---|---|---|---|---|
| all trials | $\beta_0$ 0.16<br>$\beta_1$ 0.38$^{***}$ | $\beta_0$ 0.18<br>$\beta_1$ 0.55$^{***}$ | $\beta_0$ -0.24<br>$\beta_1$ 0.83$^{***}$ | $\beta_0$ -0.22<br>$\beta_1$ 1.08$^{***}$ | $\beta_0$ -0.06<br>$\beta_1$ 0.40$^{***}$ | $\beta_0$ 0.05<br>$\beta_1$ 0.37$^{***}$ | $\beta_0$ 0.10<br>$\beta_1$ 7.68$^{***}$ |
| $EV_L > EV_R$ | $\beta_0$ 0.10<br>$\beta_1$ 0.18$^{***}$ | $\beta_0$ 0.10<br>$\beta_1$ 0.25$^{***}$ | $\beta_0$ -0.11<br>$\beta_1$ 0.37$^{***}$ | $\beta_0$ -0.14<br>$\beta_1$ 0.77$^{***}$ | $\beta_0$ 0.09<br>$\beta_1$ 0.34$^{***}$ | $\beta_0$ 0.11<br>$\beta_1$ 0.48$^{***}$ | $\beta_0$ 0.20<br>$\beta_1$ 7.94$^{***}$ |
| $EV_R > EV_L$ | $\beta_0$ 0.20<br>$\beta_1$ 0.59$^{***}$ | $\beta_0$ 0.25<br>$\beta_1$ 0.85$^{***}$ | $\beta_0$ -0.31<br>$\beta_1$ 1.26$^{***}$ | $\beta_0$ -0.31<br>$\beta_1$ 1.36$^{***}$ | $\beta_0$ 0.02<br>$\beta_1$ 0.46$^{***}$ | $\beta_0$ 0.00<br>$\beta_1$ 0.26$^{***}$ | $\beta_0$ -0.03<br>$\beta_1$ 7.52$^{***}$ |

**Table ST5. Weights for the logistic regression of choice and difference in looking time for the two screen sides.** Regression weights ($\beta_1$) and significance (***p<0.001) for results in Fig. 2B.

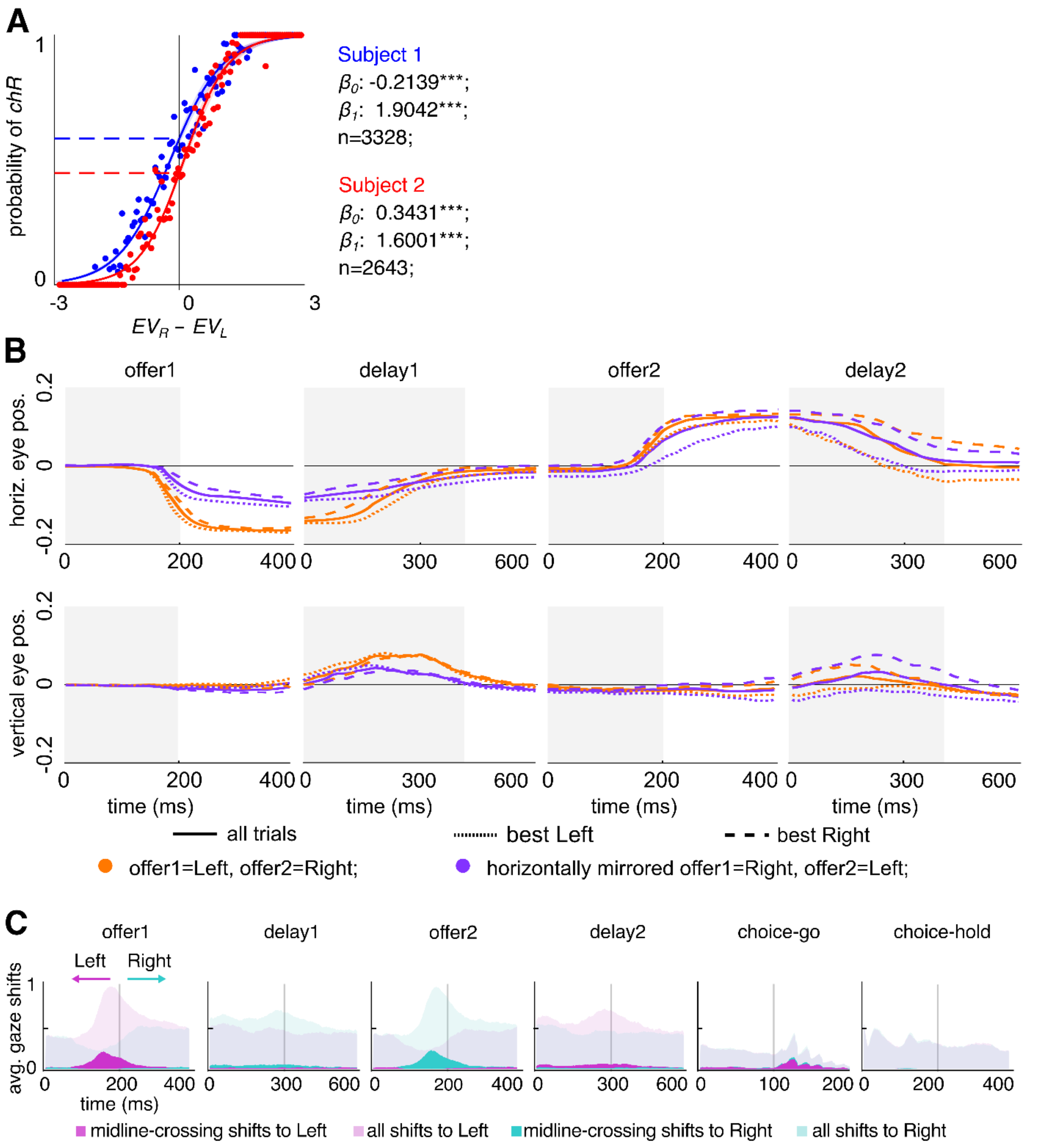


**Supplementary Figure S1. A.** Psychometric curves. Fits are made by using a logistic regression model of choice vs difference in expected values as in Fig.1B but for each subject. **B.** Top: median of horizontal eye position during task time execution (note: data for trials with first offer presented on the right screen side, blue lines, are horizontally mirrored); bottom: median of vertical eye position during task time. Solid lines include all trials, dotted lines only include trials where $EV_L > EV_R$, dashed lines only include trials where $EV_R > EV_L$. Data are combined from all trials in both subjects, in all experimental sessions. In all panels, data for the two orders of offer presentation are shown in green if first offer is presented on the left screen side, or blue if first offer is presented on the right side. **C.** Time histogram of gaze shifts detected as monotonic drifts in horizontal eye coordinate with duration at least 25ms. Midline crossing shifts are reported by solid areas.

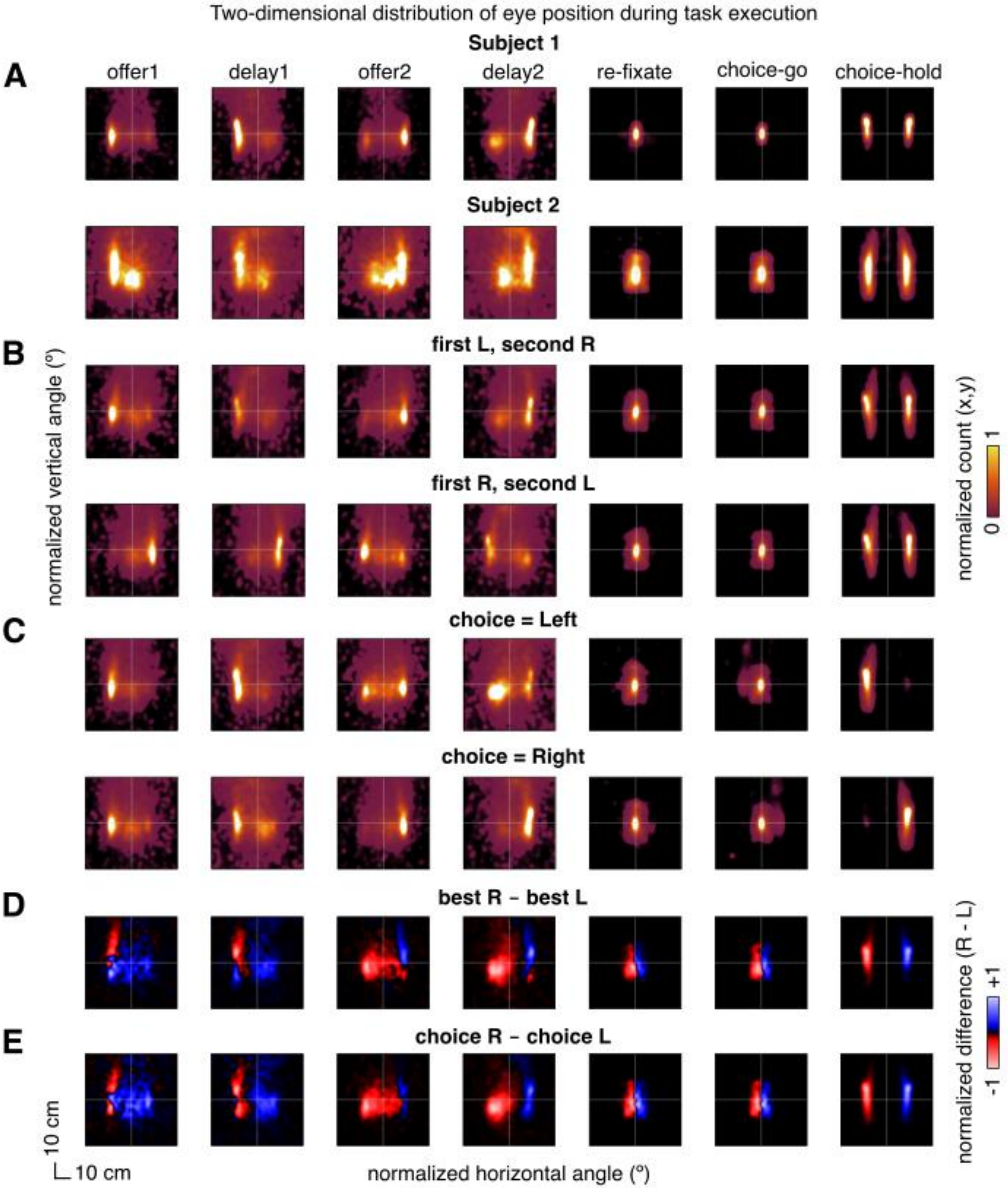

Two-dimensional distribution of eye position during task execution
Subject 1
A
offer1
delay1
offer2
delay2
re-fixate
choice-go
choice-hold
Subject 2
B
first L, second R
first R, second L
C
choice = Left
choice = Right
D
best R - best L
E
choice R - choice L
normalized vertical angle (°)
normalized horizontal angle (°)
normalized count (x,y)
0
1
normalized difference (R - L)
-1
+1
10 cm
10 cm

**Supplementary Figure S2. A.** Distribution of eye position during all task epochs. Panels in the top row include all trials across sessions for subject 1, Panels in the bottom row include trials from sessions from subject 2. **B.** Same as A, but panels in the top row include data for all sessions and for both subjects for trials with first offer presented on the left screen side (second offer on right side), while panels in the bottom row include trials with first offer on the right screen side (second offer on left side). **C.** Same as B, but panels in the top row include all trials across sessions in both subjects where the offer on the left screen side was chosen, while panels in the bottom row include trials with choice for the offer on the right screen side. **D.** Difference between the distribution of eye position with best offer is on the right side (blue, top in C) and best offer is on the left side (green, bottom in C). **E.** Difference between the distribution of eye position with choice for the right offer (blue, top in D) and choice for the left offer (green, bottom in D).

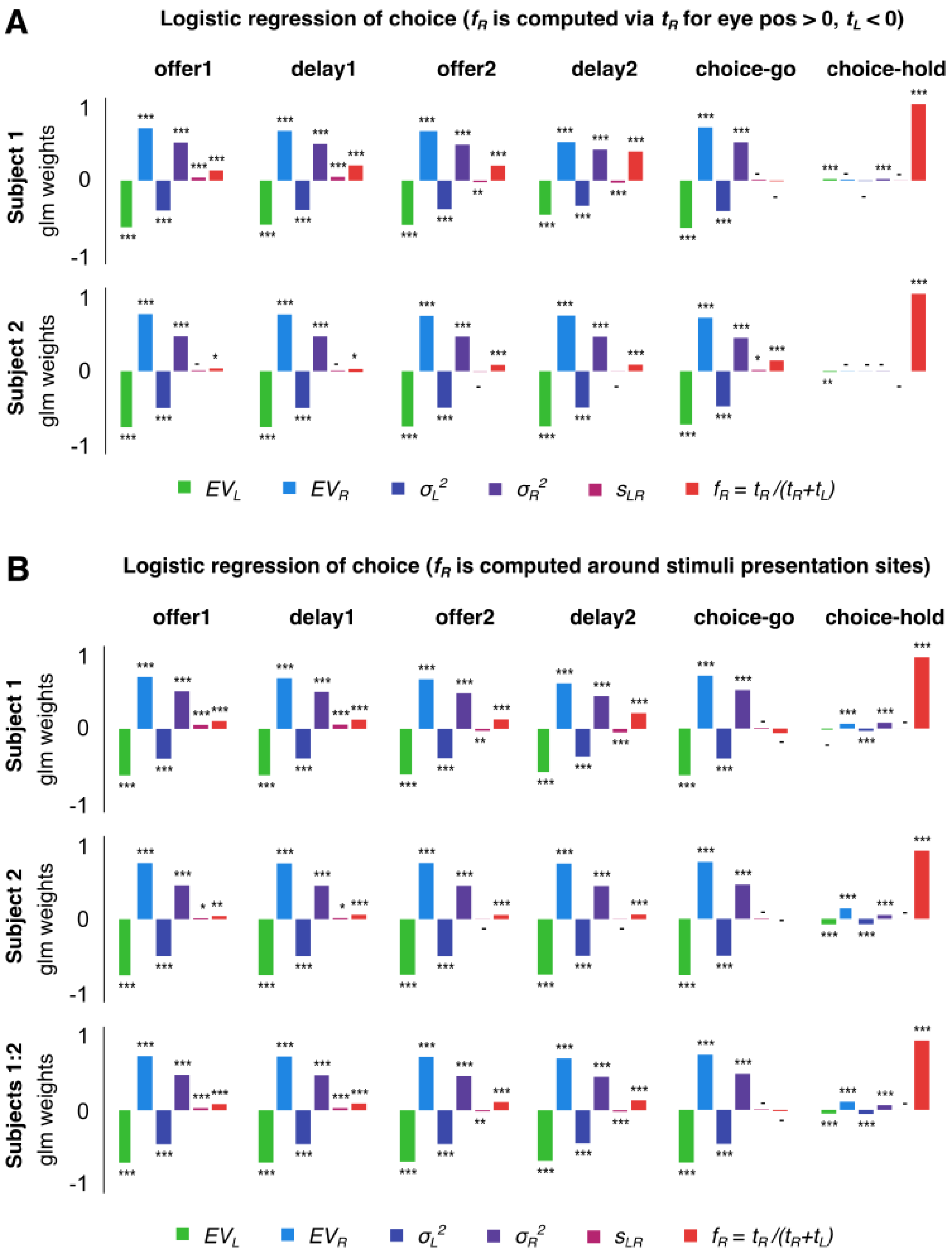


**Supplementary Figure S3. A.** Same as in Fig. 2C but separated by subjects (top: subject 1; bottom: subject 2). **B.** Same as A but fraction of time refers to time spent on L/R physical stimuli presentation sites rather L/R screen sides. Left presentation site: horizontal coordinates between -12.2±1.9cm; right site: +12.2±1.9cm.

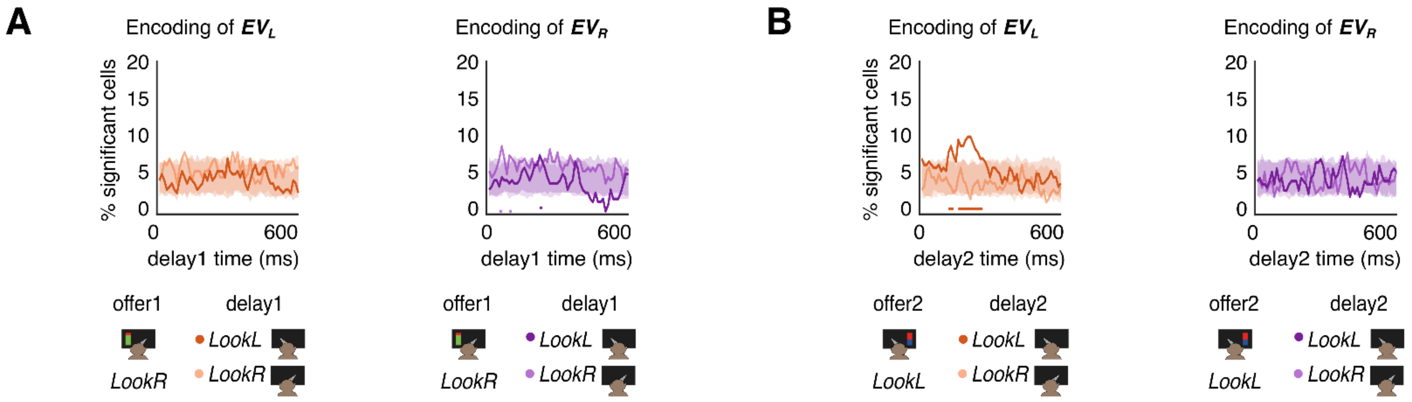


**Supplementary Figure S4. A.** Same as Fig. 3F, but *offer1 LookR*. **B.** Same as Fig. 3G, but *offer2 LookL*.

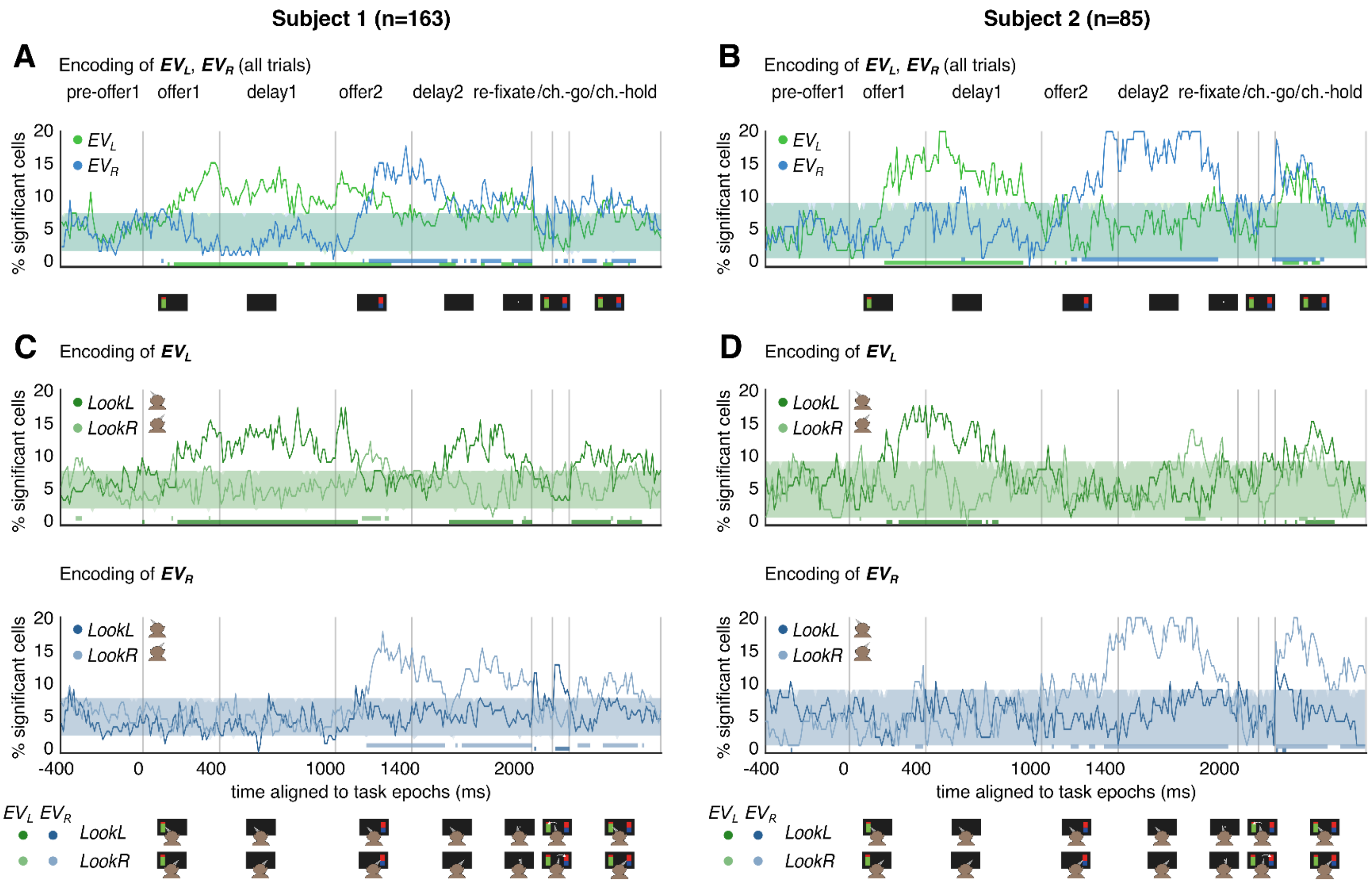


**Supplementary Figure S5. A.** Same as Fig. 3B for subject 1. **B.** Same as Fig. 3D for subject 2. **C.** Same as Fig. 3B for subject 2. **D.** Same as Fig. 3D for subject 2.

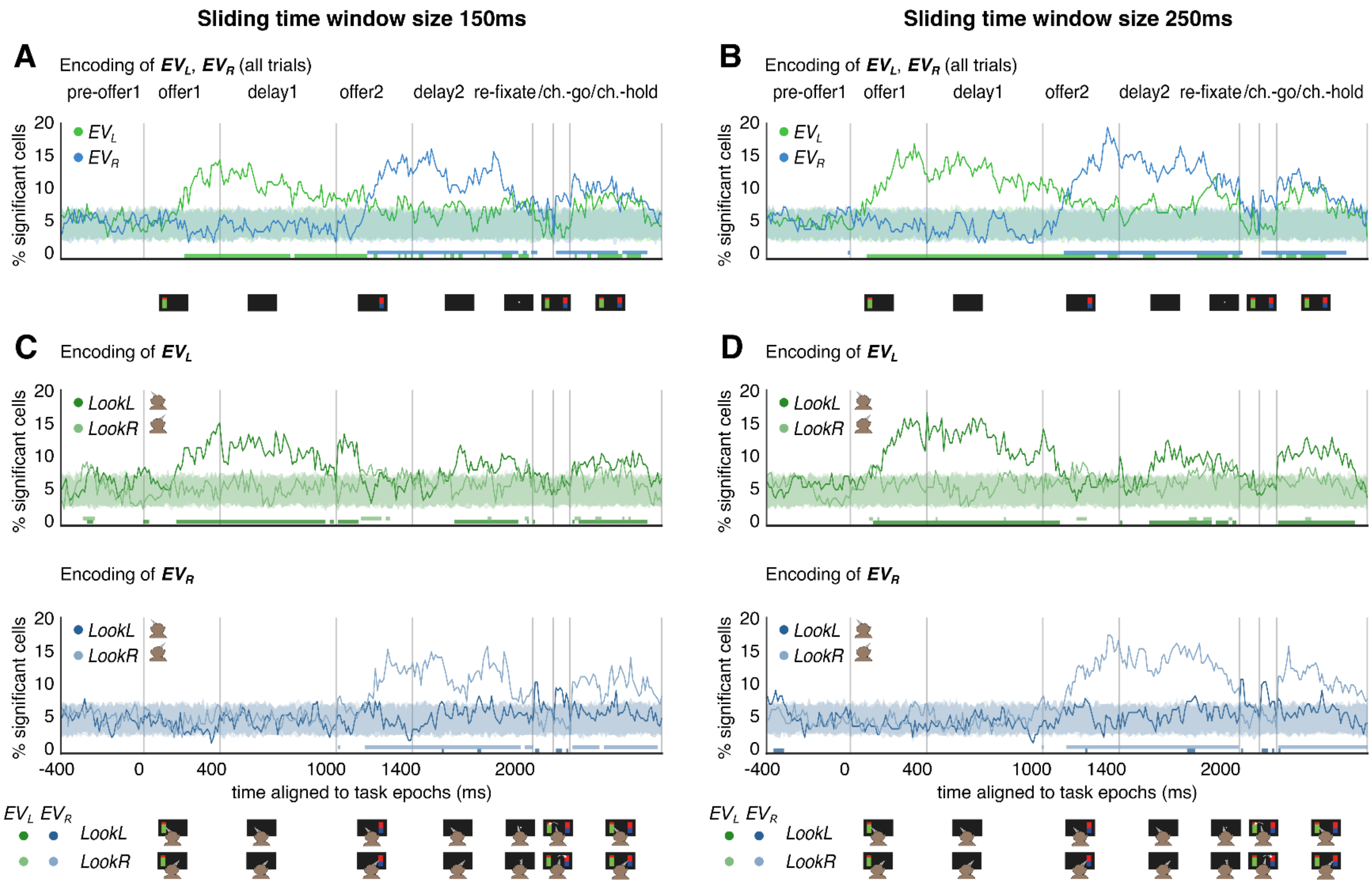


**Supplementary Figure S6. A, C.** Same as Fig. 3 B, D but for sliding time windows of size 150 ms. **B, D.** Same as Fig. 3 B, D. but for sliding time windows of size 250 ms.

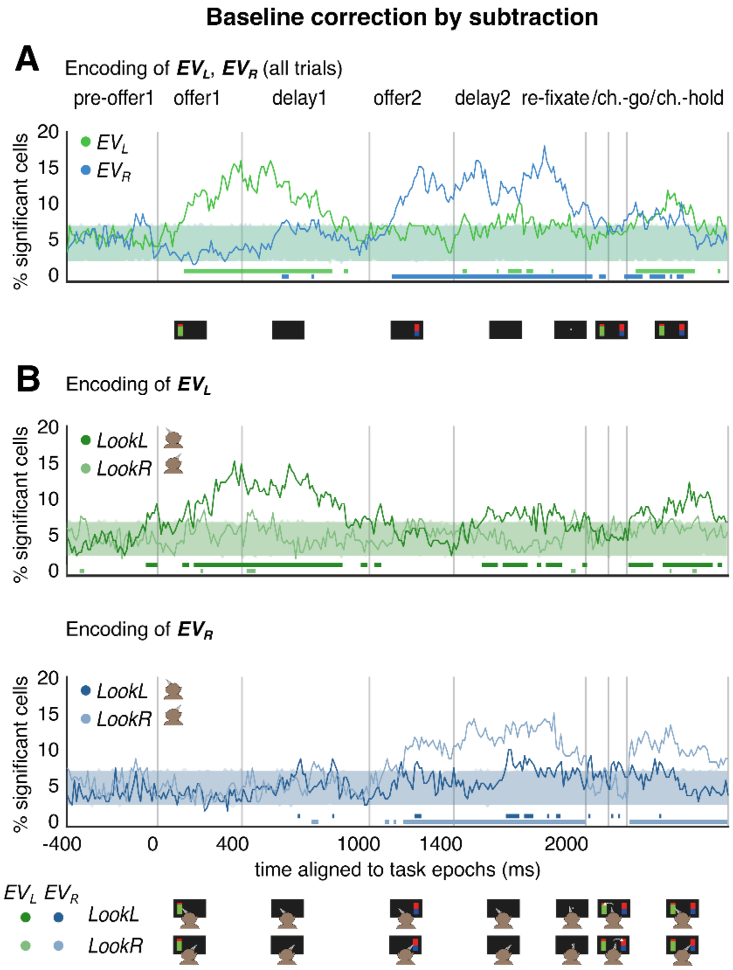


**Supplementary Figure S7. A.** Same as Fig. 3B but for baseline correction by subtraction of spike rate in each time bin by time-averaged spike rate in *pre-offer1* epoch time. **B.** Same as Fig. 3D but for baseline correction as in A.

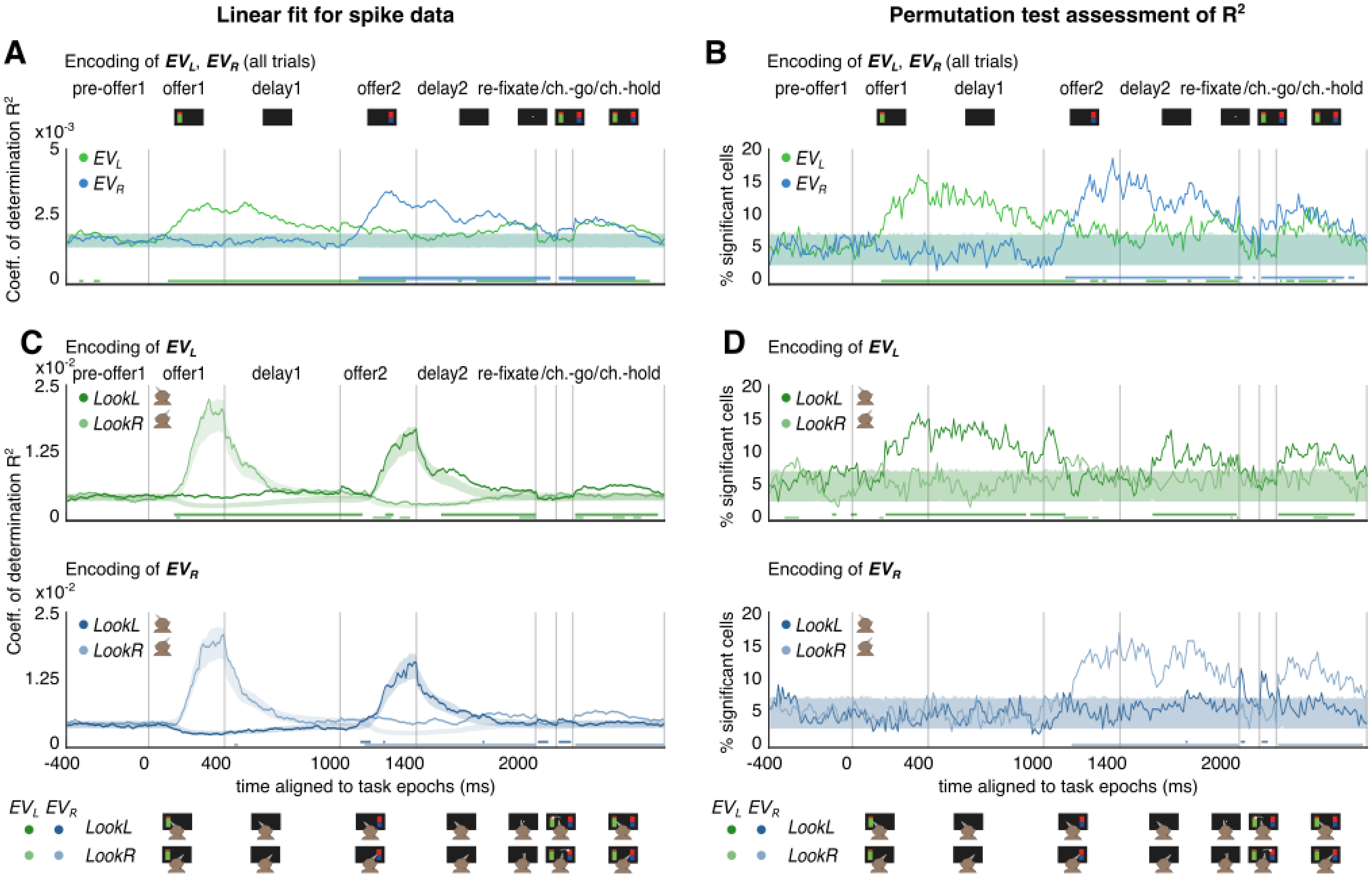


**Supplementary Figure S8. A.** Coefficient of determination $R^2$ for the results in Fig. 3B. Top: $R^2$ for the linear encoding of $EV_L$ (green) and $EV_R$ (blue) throughout the trial. Shaded areas report 5-to-95th percentile of $R^2$ distribution computed by n=1000 independent shuffles of the trial order for both EVs and spike rates. Colored lines at the bottom of the panel show time bins with $R^2$ above 95th percentile (run length assessment). **B.** Assessment of the significance of $R^2$ in panel A via permutation tests. **C.** Same as A but focusing on $EV_L$ (top) and $EV_R$ (bottom), comparing in each time bin results for trials where subjects mostly *LookL* (green or blue, respectively) vs *LookR* (light green or light blue, respectively). **D.** Same as C, but for results in panel B.

#### Sub-sampling to even trial size

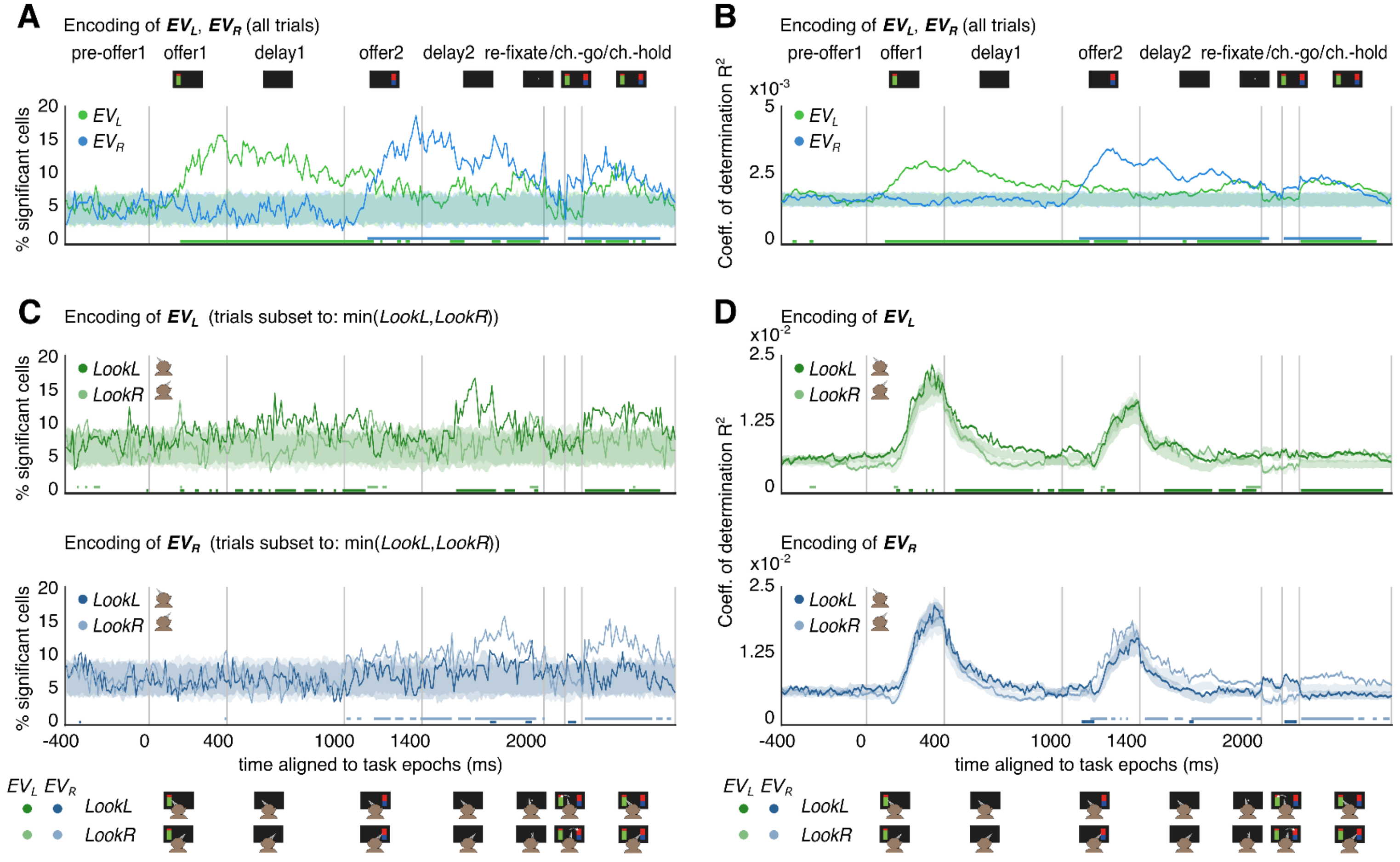


**Supplementary Figure S9. A.** Same as in Fig. 3B. **B.** Same as Supp. Fig. S8A. **C.** Same as Fig 3D, but for sub-sampled trial pools of even size. In each time bin, the number of trials for both *LookL* and *LookR* are matched in each session, and they are set to $n(t) = \min$ (number of trials *LookL*, number of trials *LookR*). The number of subsets at each time bin is given by $m(t) = [N/n(t)]$, with $N$ the total number of trials in each session. The average fraction of significant cells is computed for each subset, then fractions for all subsets are averaged. **D.** Average coefficients of determination $R^2$ for the sub-sampled results in C (to compare with Supp. Fig. S8C). Overall, sub-sampled results qualitatively match main results in Fig. 3, but they are weaker due to the lower number of trials used.

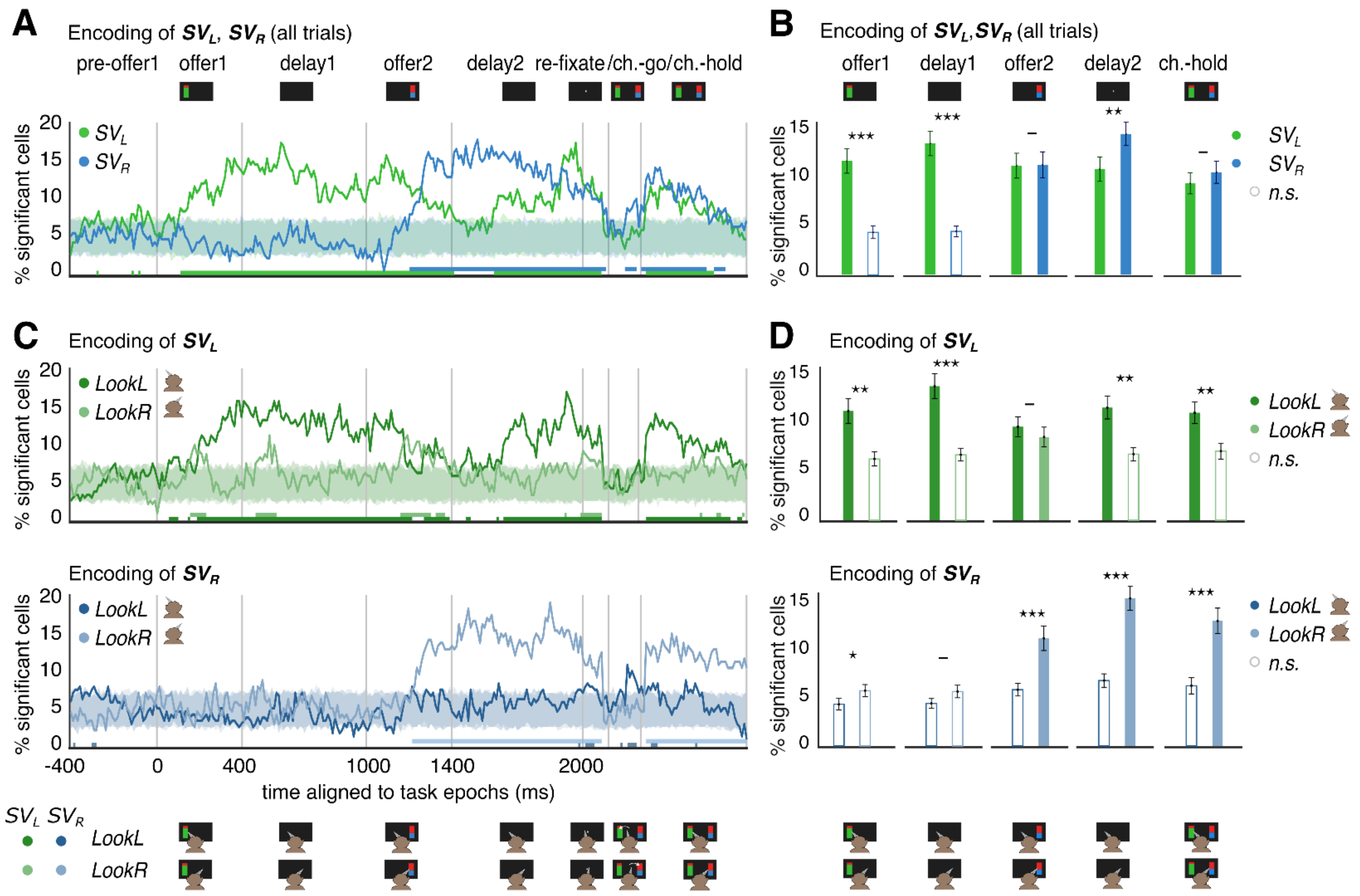


**Supplementary Figure S10. A.** Same as Fig. 3B but for Subjective Value ($SV = w_1 EV + w_2 \sigma^2$; $w_1, w_2$ previously computed in each session via logistic regression of the choice: $\text{logit}(chR) = w_0 + w_1 EV + w_2 \sigma^2$). **B.** Same as Fig. 3C but for $SV$s. **C.** Same as Fig. 3D but for $SV$s. **D.** Same as Fig. 3E but for $SV$s.

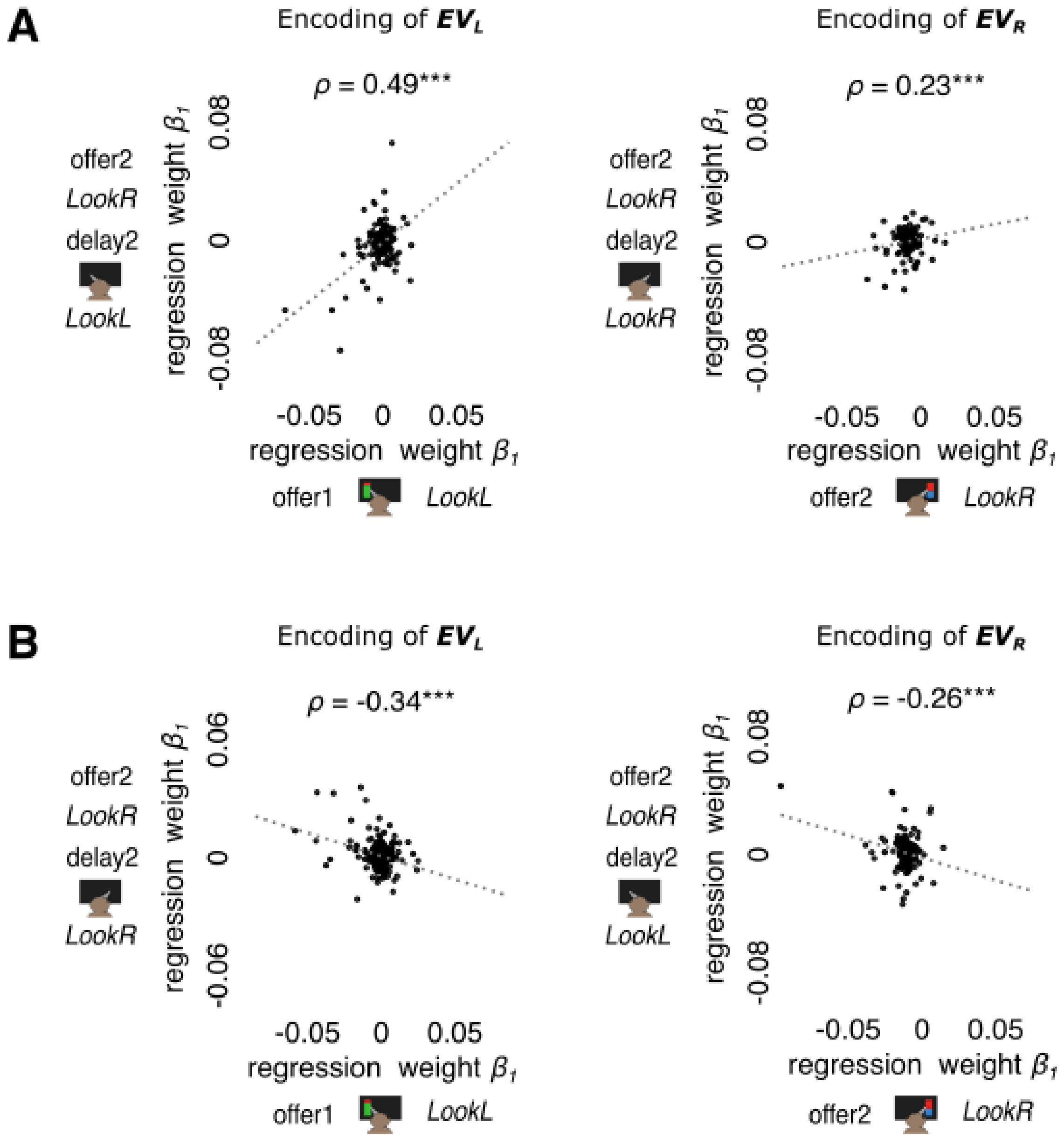


**Supplementary Figure S11. A.** Correlation of regression weights $\beta_1$ for offer $EV$s and the average firing rate ($\eta = \beta_0 + \beta_1 EV$) in ipsilateral looking conditions for *offer 1* (left, x-axis), *offer 2* (right, x-axis) and *delay 2* (y-axis). We considered spiking activity and *LookL/LookR* (as negative/positive average eye position) in the last 200 ms of all analyzed epoch times. **B.** Same as A, but for contralateral looking conditions for *offer 1* (left, x-axis), *offer 2* (right, x-axis) and *delay 2* (y-axis).
